# Switch-TRIBE: Lineage-specific high-resolution profiling of endogenous RNA-protein interactions

**DOI:** 10.64898/2026.07.29.741501

**Authors:** Yichao Hu, Alireza Khademi, Xiaohang Yang, Howard D. Lipshitz

## Abstract

RNA-binding proteins (RBPs) play vital roles in regulation of cell fate and function. Here, we introduce Switch-TRIBE, which conditionally tags endogenous RBPs with the adenosine deaminase catalytic domain to detect physiological RBP-RNA interactions. We apply Switch-TRIBE to the TRIM-NHL RBPs, BRAT and MEI-P26, essential regulators of Drosophila neurogenesis. Single-cell Switch-TRIBE in neural stem cell lineages defines cell-type specific targets and identifies novel neurogenesis regulators. Beyond target identification, Switch-TRIBE highlights the complex interplay between co-regulating RBPs, uncovering distinct binding site topographies. Our analysis uncovers a potential strategy used by BRAT and MEI-P26 to regulate lineage growth through direct binding of transcripts that encode the core metabolic and translation machineries. Switch-TRIBE provides a robust tool for characterizing RBP-RNA interactions in complex systems without compromising biological fidelity.

## INTRODUCTION

RNA-binding proteins (RBPs) dynamically interact with RNAs to orchestrate developmental programs, while aberrant binding drives diverse human pathologies (*1–3*). Consequently, precise characterization of these interactions is fundamental to understanding development and disease mechanisms. However, accurately profiling these interactions requires navigating intrinsic biological constraints. Particularly in development, RBP function can be context-dependent where regulatory networks are often restricted to specific cell types (*4*). Furthermore, RBP-RNA interactions are dose-sensitive, such that changes an RBP’s abundance has a major effect on its set of bound target RNAs (its ‘targetome’) (*1, 5*). These properties highlight the need for strategies capable of resolving cell-type-specific RBP-RNA interactions at high resolution and at endogenous RBP levels.

Standard immunoprecipitation-based approaches, including RNA immunoprecipitation (RIP) and the diverse family of crosslinking and immunoprecipitation (CLIP) methods, have been instrumental in mapping RBP targets transcriptome-wide (*6–11*). However, these workflows depend heavily on high-quality antibodies. When such reagents are unavailable, studies often resort to expressing epitope-tagged RBPs, which can introduce overexpression artifacts. Moreover, these protocols typically require large quantities of input material, limiting their capacity for cell-type-specific interrogation (*12, 13*). As a result, signals derived from bulk heterogeneous tissues inevitably obscure distinct interaction landscapes.

Emerging strategies that exploit RNA-editing signatures, such as TRIBE, HyperTRIBE, and STAMP, have substantially improved sensitivity and reduced input requirements (*14–16*). Nevertheless, these approaches typically depend on the ectopic expression of RBP-enzyme fusion proteins. Crucially, overexpression not only risks introducing non-physiological binding events but can also disturb global RNA metabolism and endogenous regulatory networks (*16*). In sensitive developmental or disease contexts, these perturbations are particularly problematic, as overexpressed or ectopic RBPs can trigger aberrant cell fate transitions or phenotypic defects, thereby confounding the biological interpretation of the results (*3*).

To bridge this methodological gap, we developed Switch-TRIBE, a strategy engineered to map cell-lineage-specific RBP-RNA interactions while preserving endogenous protein levels. We applied this approach to investigate the Drosophila TRIM-NHL proteins, BRAT and MEI-P26, essential regulators of neurogenesis whose specific RNA targets have remained elusive due to the challenges of profiling dosage-sensitive factors in complex tissues. Crucially, by coupling this strategy with single-cell sequencing (single-cell Switch-TRIBE), we resolve cell-type-specific binding profiles within the heterogeneous neuroblast lineage. Leveraging the method’s high precision to interrogate shared transcripts, we resolve distinct binding-site topographies and uncover potential competitive binding dynamics. Moreover, our analysis reveals that these paralogous RBPs employ a conserved strategy to restrict lineage growth, jointly targeting mRNAs that encode the core metabolic and translation machineries. Thus, Switch-TRIBE offers a robust and versatile framework for characterizing RBP-RNA interactions in complex systems, resolving the trade-off between sensitivity and biological fidelity.

## RESULTS

### Design of Switch-TRIBE

To enable the identification of endogenous RBP-RNA interactions with cell-lineage specificity, Switch-TRIBE employs a CRISPR-mediated knock-in strategy (**Fig. 1A**). We engineered an RBP::ADAR catalytic domain (ADARcd) fusion at the native genomic locus, separating the RBP coding sequence and the ADARcd moiety with a LoxP-flanked translational STOP cassette. In this design, the STOP cassette prevents read-through translation in the absence of Cre recombinase. To achieve precise spatiotemporal control, we utilize the binary Gal4/UAS system to drive Cre expression in defined cell lineages. Upon Cre-mediated excision of the STOP cassette, the full-length fusion protein is produced exclusively in Gal4-positive cells. By preserving endogenous cis-regulatory elements, this strategy ensures that the fusion protein is expressed at physiological levels and minimizes perturbation to native RBP functions.

**Fig. 1:**
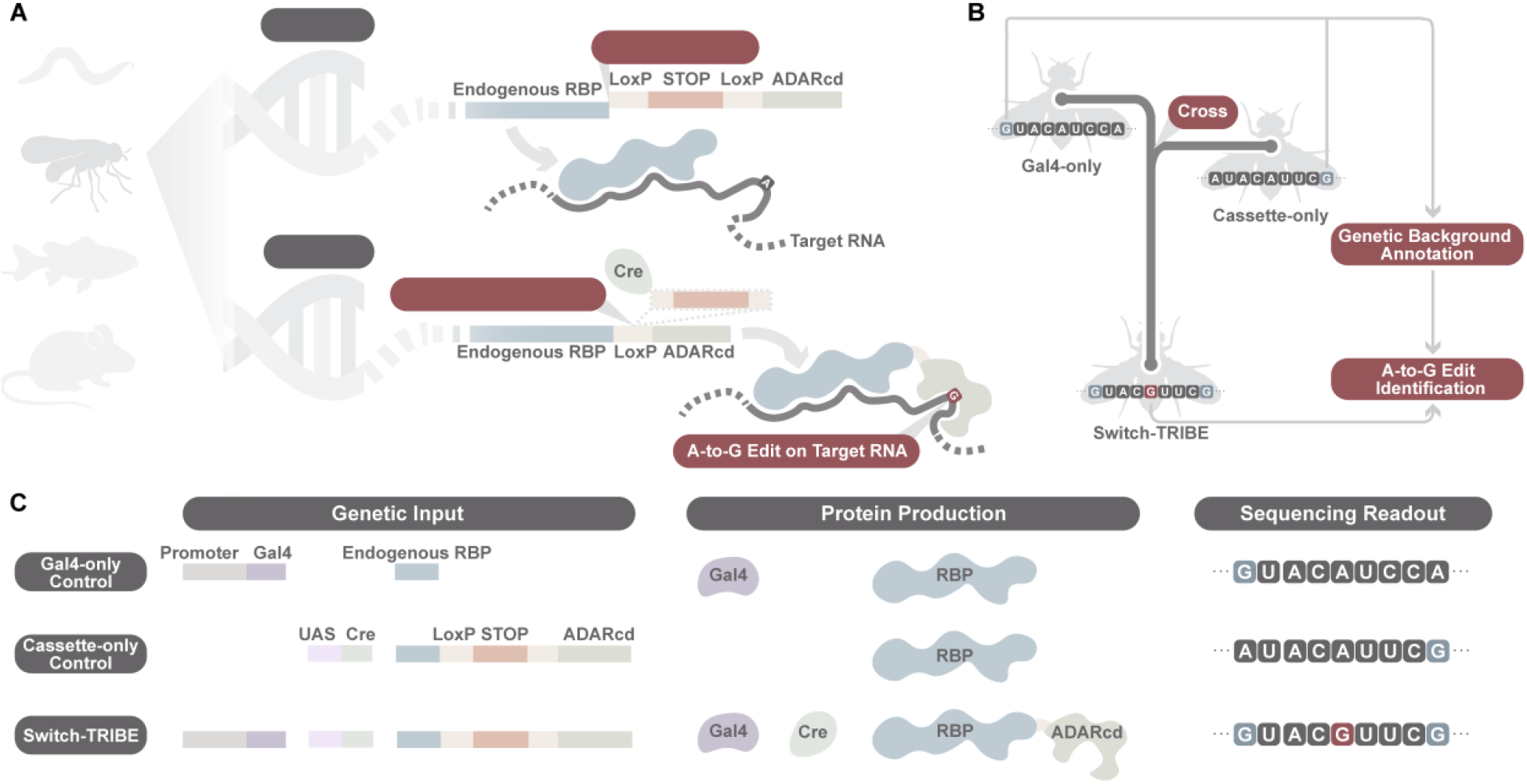
Overview of Switch-TRIBE design. **(A)** Schematic of the genomic organization and activation logic of Switch-TRIBE. The endogenous RBP locus is engineered via CRISPR knock-in to contain a LoxP-STOP-LoxP-ADARcd cassette. In the absence of Cre recombinase, the STOP cassette introduces an upstream termination signal, keeping the RBP::ADARcd fusion in an “OFF” state in most cells. In the cell types of interest, tissue-specific expression of Cre mediates excision of the STOP cassette, thereby switching the fusion to an “ON” state and enabling translation of the RBP::ADARcd protein under endogenous cis-regulatory control. The resulting fusion protein introduces A-to-I edits on bound RNAs, which are detected as A-to-G substitutions by high-throughput sequencing (*16*). Cre recombinase is driven by the Gal4-UAS system, allowing modular activation of Switch-TRIBE simply by exchanging Gal4 driver strains. **(B)** Strategy for distinguishing genuine Switch-TRIBE edits from inherited polymorphisms. Parent strains used to generate the Switch-TRIBE progeny, including the Gal4 driver line and the line carrying the Switch-TRIBE cassette, serve as baseline references for annotating background SNPs. Observed A-to-G mismatches in the progeny fall into two categories: genetic SNPs (blue) or Switch-TRIBE-dependent edits (red). By comparing A-to-G events in the progeny with those present in each parental genome, genuine RBP::ADARcd-catalyzed edits can be reliably identified. **(C)** Summary of genetic components, expected protein expression states, and sequencing outcomes for Switch-TRIBE and control conditions. The Gal4-only control contains a tissue-specific Gal4 driver but lacks the Switch-TRIBE cassette. The cassette-only control carries the CRISPR knock-in LoxP-STOP-LoxP-ADARcd cassette and the UAS-Cre transgene, which remain transcriptionally silent in the absence of Gal4. Together, these controls allow systematic assessment of background SNPs, off-target editing, and Cre-dependent activation.

Distinguishing authentic RNA edits from genomic variants can be confounded by genetic heterogeneity, particularly where SNP annotations are incomplete (*13*). To resolve this ambiguity and eliminate false positives arising from inherited polymorphisms, we implemented a rigorous genetic control strategy (**Fig. 1B**). Experimental progeny are generated by crossing a specific Gal4 driver strain with the Switch-TRIBE knock-in strain. Crucially, neither parent alone produces the active RBP::ADARcd fusion, as the knock-in parent retains the STOP cassette and the driver parent lacks the fusion allele entirely (**Fig. 1C**). By carrying out RNA-seq on both parental lines together with the Switch-TRIBE progeny, we are able to accurately subtract genomic variants and ensure the robust identification of genuine RBP-dependent editing sites.

### Switch-TRIBE enables robust and high-fidelity identification of RBP targets in complex tissues

To establish Switch-TRIBE as a generalizable platform for mapping RBP-RNA interactions in developing tissues, we selected the Drosophila TRIM-NHL protein BRAT, a critical determinant of neuroblast lineage differentiation, loss of which drives malignant tumor formation in the larval brain (*17–19*). Although several downstream effectors involved in neuroblast tumor initiation have been described, technical limitations have precluded the global identification of direct RNA targets. Consequently, a genome-wide map of RNAs directly bound by BRAT in the brain remains absent, with the notable exception of the *zelda* (*zld)* mRNA (*20*).

We engineered a BRAT::ADARcd Switch-TRIBE line and induced its expression using the insc-Gal4 driver, which is expressed in larval neuroblast lineages (**Fig. 2A, Fig. S1A**).

**Fig. 2:**
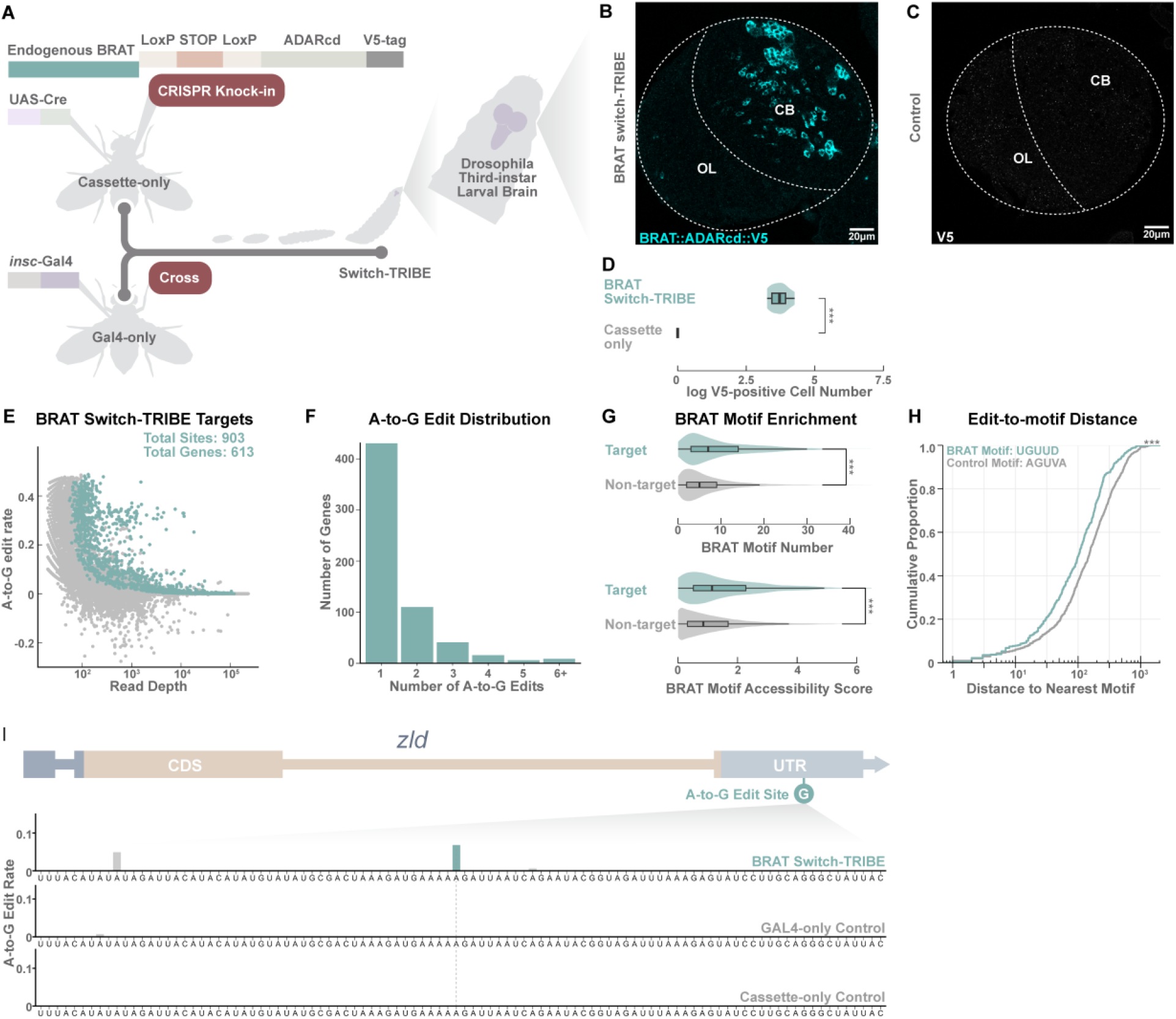
Switch-TRIBE profiling of BRAT targetomes during Drosophila neurogenesis. **(A)** Schematic of the BRAT Switch-TRIBE experimental workflow. A Drosophila strain harboring the *brat* knock-in allele (LoxP-STOP-LoxP-ADARcd-V5) was crossed with the *insc*-Gal4 driver to induce expression specifically in larval neuroblast lineages. Brains from third-instar larvae were dissected for high-throughput RNA sequencing. **(B)** Immunofluorescence validation of fusion protein expression. In Switch-TRIBE progeny, *insc*-Gal4 drives Cre-mediated activation specifically in central brain neuroblast lineages (outlined). Staining for the V5 epitope confirms robust production of the BRAT::ADARcd fusion. CB: central brain. OL, optic lobe. **(C)** Control for leakage specificity. Larvae lacking the *insc*-Gal4 driver show no detectable V5 signal, confirming that production of BRAT::ADARcd is strictly dependent on Gal4 activation. **(D)** Quantification of V5-positive cells. Bars represent cell counts per sample on a logarithmic scale. Significance was assessed using the Wilcoxon rank-sum test (*** *P* < 0.001). **(E)** Scatter plot displaying A-to-G editing sites detected in BRAT Switch-TRIBE samples. Sites with editing rates between 0 and 0.5 that passed statistical significance thresholds are classified as BRAT Switch-TRIBE targets (cyan), while non-significant sites are shown in grey. **(F)** Distribution of editing site frequency per gene among identified BRAT target transcripts. **(G)** Enrichment of BRAT binding motifs. Top: BRAT target transcripts contain significantly more BRAT motifs than co-expressed non-targets. Bottom: Cumulative motif accessibility scores are elevated in BRAT targets. Significance was determined using the Wilcoxon rank-sum test (*** *P* < 0.001). **(H)** Spatial proximity of editing sites to motifs. Cumulative distribution curves show the distance from editing sites to the nearest BRAT motif (UGUUD, cyan) compared to a control motif (AGUVA, grey). *P* values derived from the Kolmogorov-Smirnov test (*D* = 0.168, *P* = 6.6 × 10⁻⁸). **(I)** Example of BRAT Switch-TRIBE A-to-G edits on the known BRAT target *zld* (*20*). Bars show A-to-G editing rates for Switch-TRIBE samples, Gal4-only controls, and cassette-only controls (mean of n = 3 biological replicates per genotype). Significant sites are colored cyan; non-significant sites are grey.

Immunostaining confirmed robust induction of the fusion protein specifically restricted to central brain neuroblast lineages (**Fig. 2B,D**) without any adverse effect on neurodevelopment **(Fig. S1C**). Crucially, the cassette-only control (lacking the Gal4 driver) showed no detectable fusion protein, confirming the strict Gal4-dependence of fusion-protein expression (**Fig. 2C,D**).

To systematically distinguish authentic editing events, we implemented a rigorous strategy by profiling three biological replicates each of Gal4-only and cassette-only controls to annotate background mismatches arising from genetic variation. High-confidence A-to-G edits in Switch-TRIBE samples were then identified (detailed in Materials and Methods).

Application of this framework to BRAT in developing third-instar larval (L3) brains yielded 903 high-confidence A-to-G editing sites distributed across transcripts from 613 target genes (**Fig. 2E,F**; targets are listed in **Table S1**). Several lines of evidence support the authenticity of these targets. First, motif analysis revealed a strong enrichment of the canonical BRAT-binding motif (UGUUD) in Switch-TRIBE target transcripts compared with co-expressed non-targets (**Fig. 2G**). Furthermore, target transcripts exhibited significantly elevated cumulative motif accessibility scores, consistent with preferential binding by BRAT (*21, 22*) (**Fig. 2G**). We further verified these findings through spatial analysis of editing sites. Cumulative distribution analysis demonstrated that Switch-TRIBE edits were significantly closer to predicted BRAT motifs than to control sequences, reinforcing the link between RBP binding and adenosine deamination (**Fig. 2H**). Moreover, the identified target set showed significant overlap with our previous BRAT RIP-seq data from early embryos (*22*) despite the distinct developmental contexts (*P* = 4.48 x 10^-8^). Notably, Switch-TRIBE successfully captured editing events on *zld* transcripts, thereby validating our method by identification of a direct BRAT target previously identified in embryos (*22*) and L3 brains (*20*) (**Fig. 2I**).

To evaluate generalizability, we extended the approach to a second TRIM-NHL family member, MEI-P26, in the developing CNS (**Fig. 3A-C, Fig. S1B**). As for the BRAT::ADARcd Switch-TRIBE line, MEI-P26::ADARcd Switch-TRIBE expression after Insc-Gal4 excision of the STOP cassette was restricted to the central brain neuroblast lineage (**Fig. 3B,C**) without any adverse effect on neurodevelopment (**Fig. S1C**). MEI-P26 Switch-TRIBE identified a set of targets that exhibited enrichment of a previously identified binding motif (*21*) (**Fig. 3D-G, Table S1**) and a highly significant overlap (*P* = 2.62 x 10^-62^) between our Switch-TRIBE-identified targets and a set of targets previously identified in Drosophila S2 cells using iCLIP (*23*), including the *Hrb27c* mRNA (**Fig. 3H**).

**Fig. 3:**
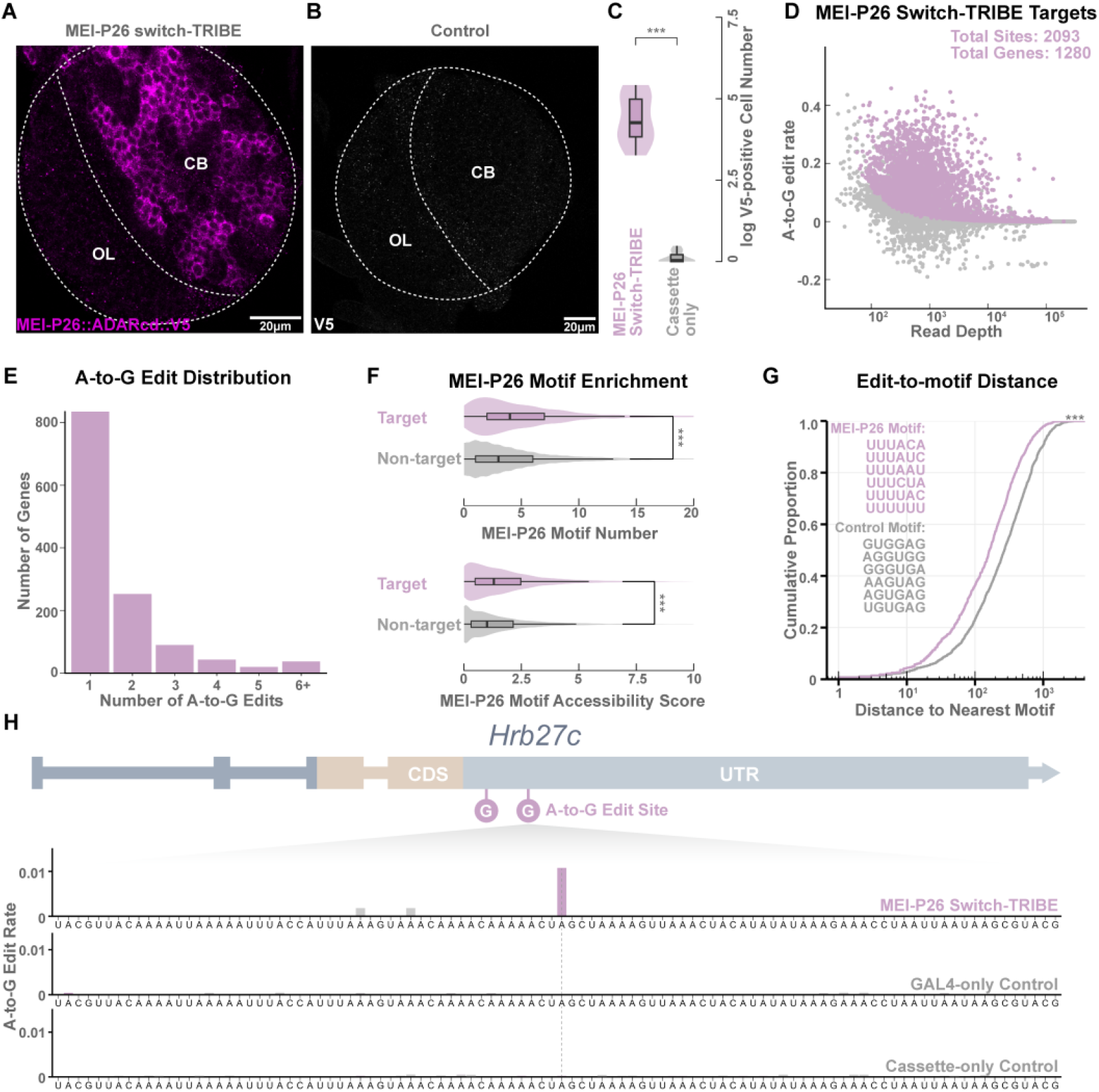
Switch-TRIBE profiling of MEI-P26 RNA targetomes during Drosophila neurogenesis. **(A)** Immunofluorescence validation of fusion protein expression. In Switch-TRIBE progeny, *insc*-Gal4 drives Cre-mediated activation specifically in central brain neuroblast lineages (outlined). Staining for the V5 epitope confirms robust production of the MEI-P26::ADARcd fusion. CB: central brain. OL, optic lobe. **(B)** Control for leakage specificity. Larvae lacking the *insc*-Gal4 driver show no detectable V5 signal, confirming that production of MEI-P26::ADARcd is strictly dependent on Gal4 activation. **(C)** Quantification of V5-positive cells. Bars represent cell counts per sample on a logarithmic scale. Significance was assessed using the Wilcoxon rank-sum test (*** *P* < 0.001). **(D)** Scatter plot displaying A-to-G editing sites detected in MEI-P26 Switch-TRIBE samples. Sites with editing rates between 0 and 0.5 that passed statistical significance thresholds are classified as MEI-P26 Switch-TRIBE targets (magenta), while non-significant sites are shown in grey. **(E)** Distribution of editing site frequency per gene among identified MEI-P26 target transcripts. **(F)** Enrichment of MEI-P26 binding motifs. Top: MEI-P26 target transcripts contain significantly more MEI-P26 motifs than co-expressed non-targets. Bottom: Cumulative motif accessibility scores are elevated in MEI-P26 targets. Significance was determined using the Wilcoxon rank-sum test (*** *P* < 0.001). **(G)** Spatial proximity of editing sites to motifs. Cumulative distribution curves show the distance from editing sites to the nearest MEI-P26 motif (UUUACA, UUUAUC, UUUAAU, UUUCUA, UUUUAC, UUUUUU, magenta) compared to a control motif (GUGGAG, AGGUGG, GGGUGA, AAGUAG, AGUGAG, UGUGAG, grey). *P* values derived from the Kolmogorov-Smirnov test (*D* = 0.187, *** *P* < 0.001). **(H)** Example of MEI-P26 Switch-TRIBE A-to-G edits on the known MEI-P26 target *Hrb27c*(*23*). Bars show A-to-G editing rates for Switch-TRIBE samples, Gal4-only controls, and cassette-only controls (mean of *n* = 3 biological replicates per genotype). Significant sites are colored magenta; non-significant sites are grey.

Together, these results demonstrate that Switch-TRIBE enables sensitive and accurate profiling of RBP-RNA interactions in a specific lineage in a complex, developmentally dynamic tissue.

### Switch-TRIBE resolves cell-type-specific RBP-RNA interactions at single-cell resolution

RBP-RNA interactions exhibit high cell-type specificity during development (*4*). However, capturing such cell-type-restricted events is challenging for conventional IP-based methods, which average signals across heterogeneous tissues (*13*). To resolve these context-dependent regulatory landscapes, we coupled Switch-TRIBE with single-cell RNA sequencing and optimized a computational pipeline to robustly identify editing events from the inherent sparsity of the resulting single-cell transcriptomes (detailed in Materials and Methods).

We applied the single-cell Switch-TRIBE strategy separately to BRAT and MEI-P26 in L3 brains (single-cell targets are listed in **Table S1**), enabling direct benchmarking against their corresponding bulk datasets (**Fig. 4**). Given the complexity of larval brain cell types, we generated a high-resolution annotation of six major clusters derived from Type I and Type II neuroblast lineages based on differential marker expression (**Figs. S3A, S4A** as detailed in Materials and Methods). We then mapped Switch-TRIBE positive cells onto these annotated clusters (**Figs. S3B,C and S4B,C**). BRAT Switch-TRIBE positive cells were predominantly neuroblasts and ganglion mother cells that derived from Type I neuroblast lineages (**Figs. 4A, and S5A,B**), whereas MEI-P26 positive cells were primarily neuroblasts and neurons (**Figs. 4D and S5C,D**). These cellular distributions recapitulate the endogenous protein expression patterns characterized by our previous study (*24*), which we further validated by immunostaining (**Fig. 4A,D**).

**Fig. 4:**
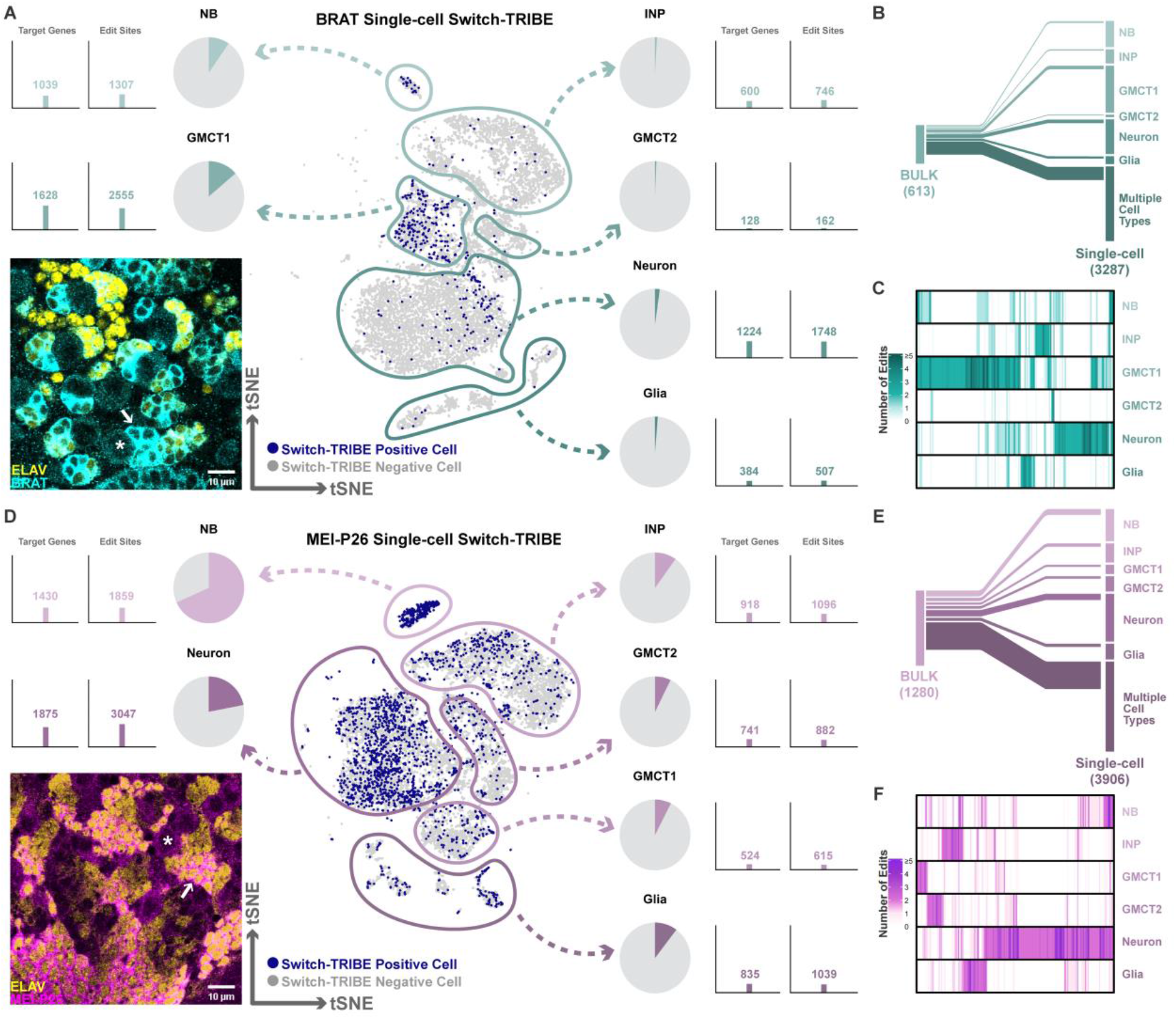
Single-cell Switch-TRIBE resolves cell-type-specific RBP-RNA targetomes during Drosophila neurogenesis. **(A)** Single-cell profiling of BRAT Switch-TRIBE activity in Drosophila third-instar larval (L3) brains. **Center:** t-SNE visualization of single-cell transcriptomes, with cells showing Switch-TRIBE activity highlighted in blue. Clusters correspond to six major neural subtypes: neuroblasts (), intermediate neural progenitors (INP), two ganglion mother cell transcriptional states (GMCT1 and GMCT2), neurons, and glia. **Top Left and Right:** Pie charts quantify the proportion of Switch-TRIBE positive cells within each cell type, while bar plots display the number of BRAT target genes and editing sites identified per cell type. **Bottom Left:** Immunostaining of BRAT and the neuronal marker ELAV in L3 central brains. Consistent with the sequencing data, BRAT is expressed most strongly in neuroblasts (large cells, asterisk) and ganglion mother cells (ELAV-negative small cells adjacent to neuroblasts, arrow) but is present at lower levels in other cell types. **(B)** Comparison of BRAT target genes identified by bulk Switch-TRIBE versus those resolved by the single-cell approach. Single-cell targets are categorized into genes edited uniquely in specific cell types or shared across multiple cell types. **(C)** Heatmap of BRAT targets identified by single-cell Switch-TRIBE. Each column represents a target gene, with the color scale indicating the total number of significant A-to-G edits detected. **(D)** Single-cell profiling of MEI-P26 Switch-TRIBE in Drosophila L3 brains. **Center:** t-SNE plot showing Switch-TRIBE positive cells (blue) across the same six neural lineages. **Top Left and Right:** Pie charts and bar plots summarize the frequency of lineage-restricted activation and the number of MEI-P26 target genes and editing sites per cell type. **Bottom Left:** Immunostaining of MEI-P26 and ELAV reveals that MEI-P26 abundance is highest in neuroblasts (large cells, asterisk) and neurons (ELAV-positive small cells, arrow). **(E)** Comparison of MEI-P26 target genes identified by bulk versus single-cell Switch-TRIBE, categorizing targets as lineage specific or shared across multiple neural cell types. **(F)** Heatmap of MEI-P26 targets identified by single-cell Switch-TRIBE. Each column corresponds to a target gene, where color intensity represents the number of significant A-to-G edits detected.

Within each of the six annotated cell types, we identified significant A-to-G editing sites and their associated target transcripts (**Fig. 4C,F**). Benchmarking against bulk Switch-TRIBE data demonstrated that the majority of bulk targets were successfully recovered, confirming the consistency of the single-cell approach (**Fig. 4B,E**). Importantly, single-cell Switch-TRIBE uncovered a substantially expanded target repertoire relative to bulk Switch-TRIBE. These results demonstrate that single-cell Switch-TRIBE can effectively resolve complex, context-dependent RBP-RNA interaction landscapes.

We next verified that Switch-TRIBE preserves endogenous transcriptional programs by comparing global gene expression profiles between Switch-TRIBE positive and negative cells within the same cell type (**Fig. S6**). The transcriptomes of these populations were highly concordant, indicating that the presence of the fusion protein does not distort global gene expression (**Fig. S6B,D**). Furthermore, the levels of *brat* and *mei-P26* transcripts showed no significant differences between Switch-TRIBE positive and negative populations in their predominant cell types (**Fig. S6A,C**). Together with the absence of any neurodevelopmental effects (**Fig. S1C**), these analyses confirm that Switch-TRIBE operates without disturbance of normal neural lineage specification and differentiation.

### Mapping convergent regulatory networks of paralogous RBPs in development

RBPs typically engage extensive target repertoires, and individual transcripts can be bound by multiple RBPs (*1*). Precise characterization of how distinct RBPs coordinate on these shared substrates requires a strategy capable of resolving interactions without introducing ectopic binding. Here we reasoned that Switch-TRIBE might provide a rigorous framework to dissect these relationships directly *in vivo*.

BRAT and MEI-P26 are structurally homologous TRIM-NHL proteins that are each essential for Drosophila neurogenesis (*17–21, 24, 25*). We first compared the molecular functions associated with their targets identified by bulk and single-cell Switch-TRIBE. Gene Ontology (GO) enrichment analyses indicated that targets of both RBPs are associated with similar functional categories (**Fig. 5A**). This resemblance further extended to their binding topographies, where A-to-G editing events for both proteins were enriched within coding sequences and 3′ UTRs (**Fig. 5B**).

**Fig. 5:**
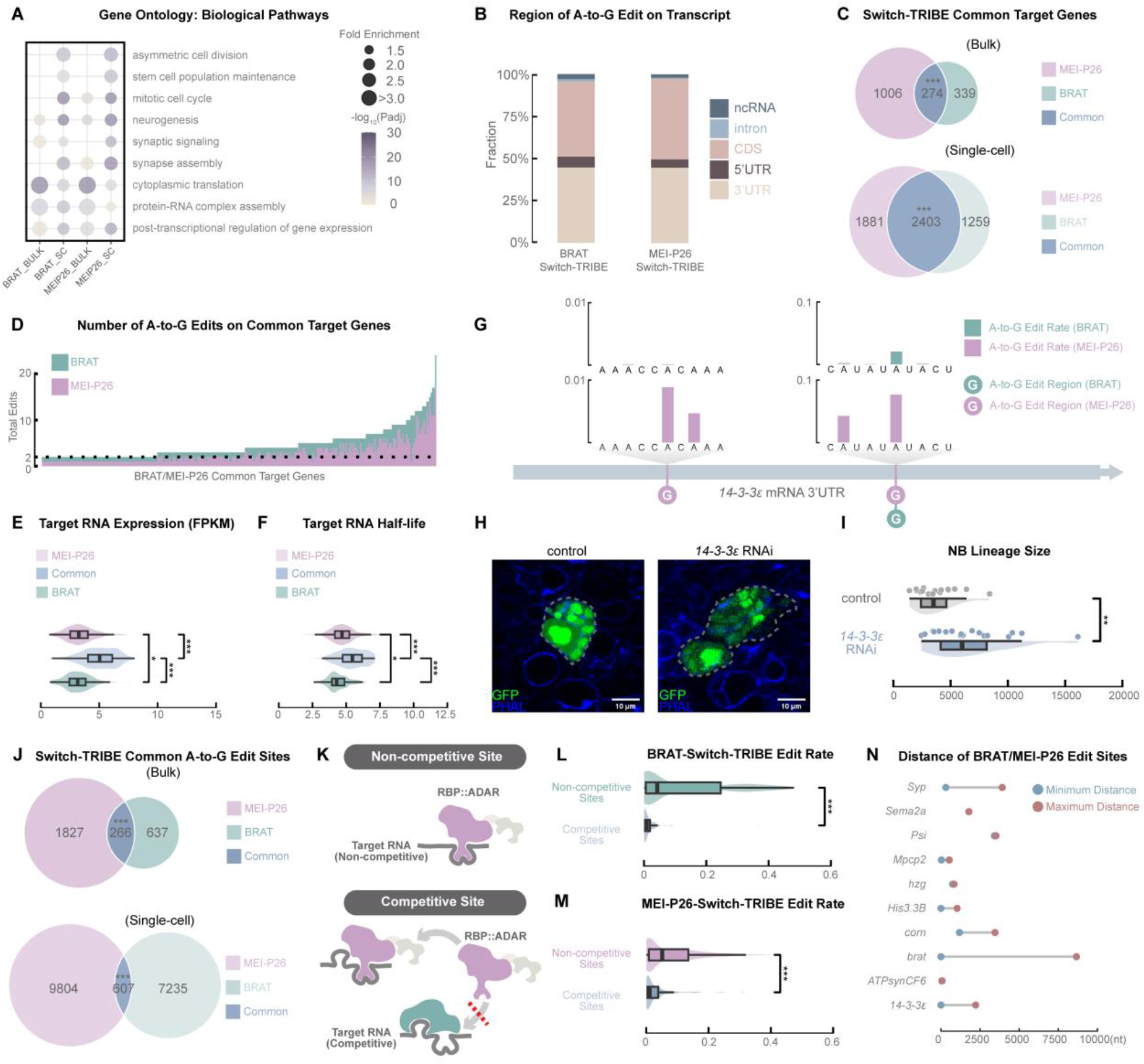
Comparative analysis reveals convergent regulation and potential competitive binding by BRAT and MEI-P26. **(A)** Functional enrichment analysis of targets identified by bulk and single-cell Switch-TRIBE. Dot plots display Gene Ontology (GO) terms of biological pathways enriched in BRAT and MEI-P26 target sets. **(B)** Transcript-wide distribution of A-to-G edits detected in bulk BRAT and bulk MEI-P26 Switch-TRIBE experiments. **(C)** Overlap of target genes identified by Switch-TRIBE. Euler diagrams compare gene sets derived from bulk (top) and single-cell (bottom) datasets. Significance was determined by the hypergeometric test (*** *P* < 0.001). **(D)** Editing site density on co-targeted genes. Columns represent individual genes targeted by both RBPs in bulk experiments, highlighting BRAT (cyan) and MEI-P26 (magenta) editing sites. **(E)** Expression levels of target transcripts. Comparison of shared targets versus BRAT-specific and MEI-P26-specific targets. Significance was determined by the Wilcoxon rank-sum test (* *P* < 0.05; *** *P* < 0.001). **(F)** RNA stability analysis comparing the half-lives of shared and RBP-specific target transcripts. RNA half-lives in third-instar larval brains were calculated using previously published dataset (*26*). Significance was determined by the Wilcoxon rank-sum test (*: P < 0.05; *** *P* < 0.001). **(G)** Phenotypic analysis of the shared target *14-3-3ε*. Immunostaining of L3 neuroblast lineages in control and *14-3-3ε* RNAi backgrounds. GFP marks Gal4-positive lineages and phalloidin labels cell boundary. **(H)** Quantification of neuroblast lineage volume. Loss of *14-3-3ε* results in larger lineage size (** *P* < 0.01, Wilcoxon rank-sum test). **(I)** Representative editing tracks on the *14-3-3ε* 3′ UTR. Bars show mean editing rates (n = 3) for BRAT (cyan) and MEI-P26 (magenta). Grey bars indicate non-significant sites. **(J)** Overlap of specific A-to-G editing sites detected in bulk (top) and single-cell (bottom) datasets. Significance was determined by the hypergeometric test (*** *P* < 0.001). **(K)** Model of competitive binding. Co-occupancy by endogenous RBPs competes with Switch-TRIBE fusion binding, potentially reducing the detectable editing rate at shared sites. **(L)** Evidence of competition in BRAT Switch-TRIBE data. Box plots compare editing rates at shared sites (competitive) versus unique sites located on shared target genes (non-competitive). Shared sites exhibit significantly lower editing rates (*** *P* < 0.001, Wilcoxon rank-sum test). **(M)** Evidence of competition in MEI-P26 Switch-TRIBE data. Comparison of editing rates for competitive versus non-competitive sites within the MEI-P26 dataset (*** *P* < 0.001, Wilcoxon rank-sum test). **(N)** Proximity of editing sites on shared targets. Dumbbell plot displays the minimum and maximum distances between BRAT and MEI-P26 editing sites on representative co-targeted transcripts, including *14-3-3ε*.

We found a substantial and statistically significant overlap between the BRAT and MEI-P26 targetomes (**Fig. 5C**). The shared targets frequently contained multiple A-to-G edits by each RBP (**Fig. 5D**). Moreover, when compared to a previously published dataset (*26*), these common targets displayed significantly higher expression levels and markedly longer RNA half-lives than targets bound by either RBP alone (**Fig. 5E,F**). These results suggest that the shared targets represent a distinct regulatory class.

To investigate the biological significance of this extensive co-targeting, we focused on shared targets enriched in neurogenesis-related GO terms (**Table S2**). Among these, we identified the *14-3-3ε* mRNA as extensively edited in its 3′ UTR by both RBPs (**Fig. 5G**); notably, we detected a shared editing site approximately 900nt 3′ of a MEI-P26-enriched editing cluster, indicating that BRAT and MEI-P26 may occupy proximal or overlapping footprints on *14-3-3ε* transcripts. Notably, editing by MEI-P26 was more extensive than by BRAT, suggesting that MEI-P26-dependent regulation might predominate. Consistent with this hypothesis, knockdown of *14-3-3ε* mRNA resulted in enlarged neuroblast lineages, similar to what we previously observed upon *mei-P26* knockdown (*24*) (**Fig. 5H,I)**. Thus, both MEI-P26 and the protein encoded by its direct target, *14-3-3ε* mRNA, function to prevent over-proliferation of neuroblast lineages.

Together, these results demonstrate proof-of-principle that Switch-TRIBE-driven, lineage-specific identification of RBP targetomes can lead to identification of new developmental effectors regulated by those RBPs.

### High-resolution comparative analysis at native stoichiometry uncovers quantitative RBP competition

Our high-resolution mapping revealed that BRAT and MEI-P26 Switch-TRIBE could catalyze edits at identical nucleotide positions on the *14-3-3ε* transcript (**Fig. 5G**). While enzymatic editing events do not strictly correspond to physical binding footprints, such precise coincidence implies that the two RBPs can occupy nearby or overlapping binding sites on the target (*14, 16*). This observation prompted us to investigate whether such coincident editing events might be widespread. Indeed, a transcriptome-wide comparison uncovered a significant overlap of A-to-G editing sites between the BRAT and MEI-P26 Switch-TRIBE datasets (**Fig. 5J**), suggesting that these paralogous TRIM-NHL RBPs frequently engage the same RNAs *in vivo*.

We hypothesized that this overlap may reflect direct competition for shared RNA binding sites, if BRAT and MEI-P26 are co-expressed in the same cell type. In a competitive binding framework, the access of an RBP::ADARcd fusion to a shared site is expected to be impeded when another endogenous RBP competes for the same motif, thereby lowering the observed editing frequency (**Fig. 5K**). To test this quantitatively, we defined ‘competitive sites’ as positions edited by both BRAT and MEI-P26, and ‘non-competitive’ sites as positions edited by only one RBP but located on shared target transcripts. Consistent with the competitive binding model, we observed significantly reduced editing rates at competitive sites (**Fig. 5L,M**). This model aligns with a previous study showing that BRAT can also recognize MEI-P26 motifs (*23*).

To further resolve the spatial organization of binding on shared transcripts, we calculated the minimum and maximum distances between BRAT and MEI-P26 editing sites for 10 common targets (**Fig. 5N**). This analysis revealed three distinct binding modes: distant binding, exemplified by the *Sema2a, corn* and *Psi* mRNAs, where BRAT and MEI-P26 editing sites occupied clearly separated regions; overlapping binding, observed in target mRNAs such as *ATPsynCF6* and *Mpcp2*, where editing sites for the two RBPs clustered in close proximity; and mixed binding, represented by the *Syp*, *brat* and *14-3-3ε* transcripts, which featured both close and distant pairs of BRAT and MEI-P26 sites (**Fig. 5N**).

To assess whether BRAT and MEI-P26 might bind their shared targets in the same or different cells within neuroblast lineages we assessed our single-cell Switch-TRIBE data for cell-type-specific edits (**Table S1**). As examples, *Syp*, *corn*, *brat*, *His3.3B* and *14-3-3ε* mRNAs were identified as co-targets of BRAT and MEI-P26 in either neuroblasts or neurons (or both) – which are cell types in which BRAT and MEI-P26 are known to be co-expressed (*24*). In contrast, *Sema2a*, *hzg* and *Psi* transcripts were not identified as co-targets in neuroblasts or neurons; thus, in these two cell types they are targeted either by BRAT or MEI-P26, but not both. *Mpcp2* and *ATPsynCF6* mRNAs were identified as co-targets in bulk but not single-cell Switch-TRIBE; thus, we cannot draw any conclusions regarding cell-type specificity of binding to these transcripts.

Together, these analyses demonstrate that Switch-TRIBE, applied at the native stoichiometry of endogenous RBPs, can resolve complex target-binding modalities.

### Endogenous profiling distinguishes direct regulation from indirect downstream consequences in neurogenesis

We next sought to apply the insights from Switch-TRIBE to previous genetic and developmental studies of BRAT and MEI-P26 in larval neuroblast lineages. The role of BRAT as a brain tumor suppressor has been well-established for more than twenty years (*17, 18*). Previous studies have identified multiple potential downstream effectors capable of rescuing the *brat* mutant tumor phenotypes upon genetic manipulation (*20, 27–31*). However, with the exception of *zld* (*20*), it has remained unclear whether these effectors are direct targets of BRAT or indirect downstream genes. Here, we categorized these effectors into direct and indirect targets using our bulk and single-cell BRAT Switch-TRIBE data (**Fig. 6A**). Notably, the previously identified effector *lncRNA:cherub* (*29*) emerged as one of the top hits in the BRAT Switch-TRIBE dataset, harboring seven A-to-G editing sites. Furthermore, nine out of 15 of these effectors were significantly upregulated in *brat* knockdown Type II neuroblast lineages (**Fig. 6A**), based on our re-analysis of a previously published dataset (*29*).

**Fig. 6:**
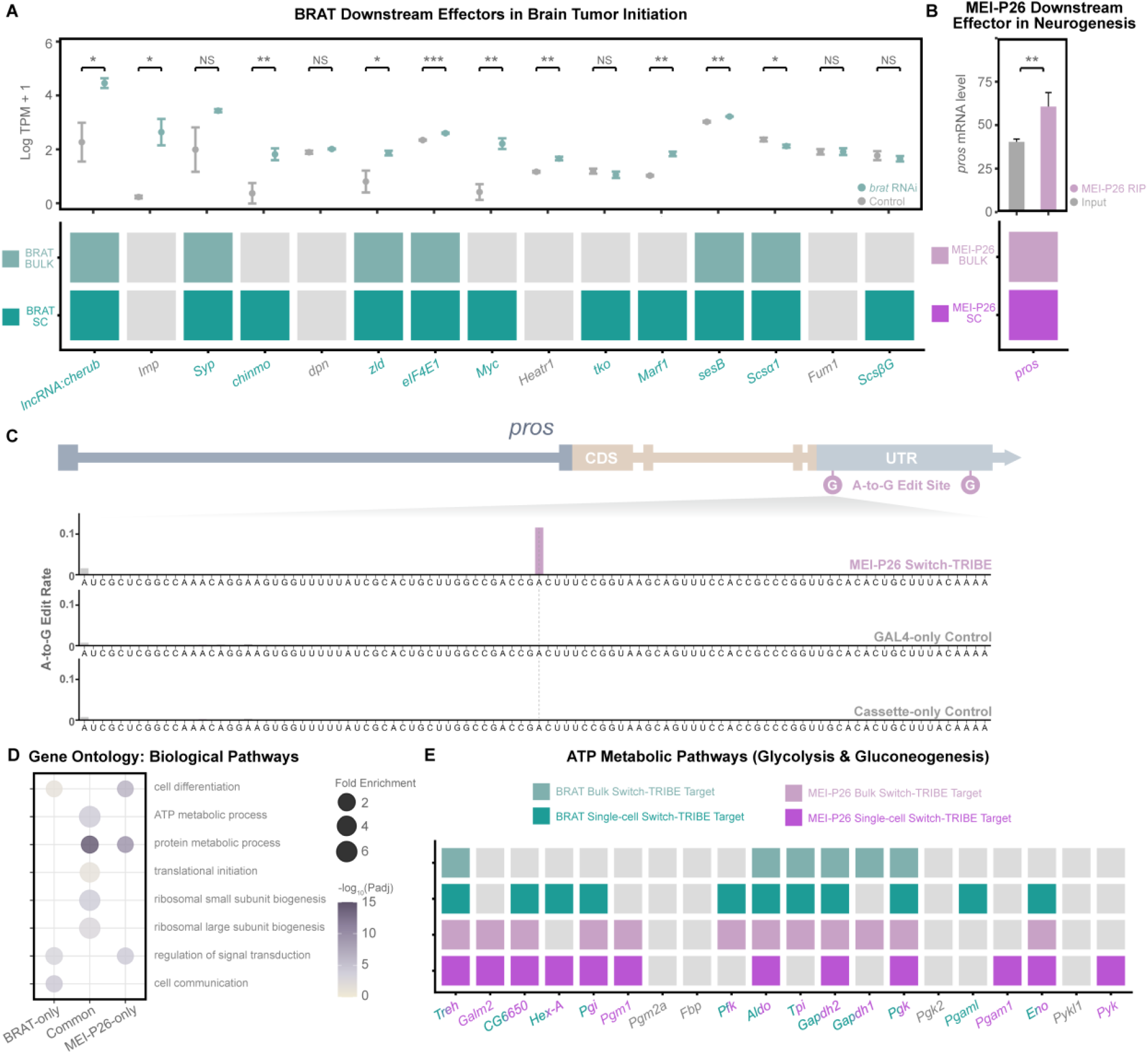
Redefining the direct regulatory landscape and metabolic impact of BRAT and MEI-P26. **(A)** Distinguishing direct targets from indirect effectors. Top: Differential expression of known BRAT downstream effectors (reported in literature to drive brain tumor initiation), quantified by RNA-seq in brat RNAi versus control type II neuroblast lineages. Data are presented as log-transformed expression levels. Significance was determined by two-tailed t-tests (* *P* < 0.05; ** *P* < 0.01; *** *P* < 0.001; NS: not significant). Bottom: Re-classification of these effectors as direct targets (validated by bulk or single-cell Switch-TRIBE) or indirect downstream genes, resolving their hierarchical position in the regulatory network. **(B)** Validation of *pros* as a direct MEI-P26 target. Top: Enrichment of *pros* transcripts in MEI-P26 RIP-seq data. Significance was computed using DESeq2 (** adjusted *P* < 0.01). Bottom:Cross-validation of *pros* as a direct target based on reproducible identification in both bulk and single-cell MEI-P26 Switch-TRIBE datasets. **(C)** Editing profile of the pros transcript. Tracks display A-to-G editing rates in MEI-P26 Switch-TRIBE samples compared with Gal4-only and cassette-only controls (mean of n = 3 biological replicates). Significant editing sites are highlighted in magenta; non-significant sites are shown in grey. **(D)** Functional specialization and convergence. Gene Ontology (GO) enrichment analysis of BRAT-specific, MEI-P26-specific, and common target genes. Dot size represents fold enrichment, and color scale indicates adjusted P values. **(E)** Map of direct targets within central metabolic pathways. Schematic of the glycolysis and gluconeogenesis pathways; genes identified as direct BRAT and/or MEI-P26 targets are highlighted.

The function of MEI-P26 has been extensively studied in germline stem cell lineages (*32–34*). However, previous genetic screens have also identified it as an essential regulator of neurogenesis (*25*). We recently showed that MEI-P26 modulates neuroblast lineage size by positively regulating the transcription factor *pros* (*24*); however, in that study, whether *pros* mRNA is a direct or indirect target of MEI-P26 remained unclear. Here, our bulk and single-cell MEI-P26 Switch-TRIBE data both indicated that *pros* is a direct target of MEI-P26 in L3 brains (**Fig. 6B,C**). This finding was further validated by whole-brain RIP-seq experiments (**Fig. 6B**), where *pros* transcripts were significantly enriched in the MEI-P26 RIP compared to input.

Together, these analyses expand and extend previous findings regarding the genetic interactions downstream of these two RBPs in neurogenesis, providing a robust strategy for distinguishing direct regulation from indirect downstream consequences. This framework allows for the elucidation of regulatory networks in genetic and developmental studies. Furthermore, it facilitates the discovery of novel functional effectors by pinpointing the direct targets of RBPs.

### Switch-TRIBE reveals a conserved function for BRAT and MEI-P26 in targeting core metabolic and translation machinery

RBPs typically interact with hundreds to thousands of RNAs (*1, 2*). However, classical genetic and developmental studies have largely focused on individual downstream effectors, owing to methodological limitations (*1, 3*). Our observation that many genetically identified BRAT downstream effectors are direct BRAT targets supports a model of parallel regulation, suggesting that RBPs may orchestrate development by targeting transcripts across diverse functional pathways. We therefore interrogated the BRAT and MEI-P26 targetomes to uncover potential downstream effector pathways involved in neurogenesis.

As described above, our Switch-TRIBE data allowed us to identify an additional regulator of neurogenesis, *14-3-3ε* (**Fig. 5G-I**). Moving beyond individual targets, here we extended our investigation to define the broader downstream regulatory landscape, focusing specifically on the biological pathways enriched within the shared targetome (**Fig. 6D, Table S2**).

We found that BRAT-only and MEI-P26-only target sets were enriched for terms such as cell differentiation and regulation of signal transduction, while BRAT–MEI-P26 common targets were enriched for ATP metabolic processes and terms relevant to cytoplasmic translation (**Fig. 6D**). Previous studies have reported that genes encoding ATP metabolic enzymes are misregulated in *brat* knockdown cells (*35*); however, whether these genes are direct targets of BRAT remained unknown. Given that ATP metabolic processes are enriched in BRAT–MEI-P26 common targets rather than BRAT-only targets, we mapped both the BRAT and MEI-P26 targetomes to the glycolysis and gluconeogenesis pathways (**Fig. 6E**). Among 20 annotated genes in these pathways, we found that 16 are targets of BRAT and/or MEI-P26, and 10 are common targets. Given that upregulated metabolic activity is essential for brain tumor growth (*28*), our results are consistent with the hypothesis that BRAT and MEI-P26 may suppress neuroblast lineage overgrowth by globally targeting ATP metabolic genes.

We next focused on the term cytoplasmic translation. The BRAT–MEI-P26 common target set was enriched in GO terms including translational initiation, ribosomal small subunit biogenesis, and ribosomal large subunit biogenesis (**Fig. 6D**); indeed, the majority of genes annotated to these specific biological processes are direct targets of BRAT or MEI-P26 as identified by Switch-TRIBE (**Fig. S7**). This pattern suggests a mechanism of coordinated global translational control, where these paralogous RBPs jointly target the core machinery governing translation. At this time we do not know whether the regulatory outcome is positive or negative.

In summary, Switch-TRIBE reveals that BRAT and MEI-P26 may employ a conserved regulatory strategy to modulate neurogenesis. By uncovering that these paralogous RBPs co-regulate core metabolic and translation modules, this study demonstrates the utility of Switch-TRIBE for dissecting complex post-transcriptional networks *in vivo*.

## DISCUSSION

Here, we have introduced Switch-TRIBE as a method that profiles physiological RBP-RNA interactions with high spatiotemporal resolution. Using Drosophila neurogenesis as a paradigm, we validated the method’s fidelity and identified the targetomes of BRAT and MEI-P26. Coupling Switch-TRIBE with single-cell sequencing allowed us to dissect interaction landscapes in specific cell types within heterogeneous lineages. Crucially, the method’s high sensitivity provided a unique window into the dynamic regulatory architecture of these RBPs, uncovering binding patterns that were previously obscured by tissue heterogeneity.

A critical prerequisite for achieving such high-resolution profiling is the ability to record interaction histories with minimal disturbance to the cellular context. Recently developed methods, such as Mapit-seq (*36*) and INSCRIBE (*37*), employ antibody-tethered editing enzymes to map interactions without genetic manipulation. While these methods avoid overexpression artifacts, they require high-quality antibodies and generate edits post-fixation, thereby capturing static snapshots of the interaction landscape (*36, 37*). Conversely, genetically encoded approaches like TRIBE and STAMP function as molecular recorders, accumulating edits over time to capture transient or dynamic events (*15, 16*). However, these methods typically depend on ectopic or over-expression, which risks perturbing endogenous regulatory networks. Switch-TRIBE bridges this gap by enabling historical recording of targetomes at endogenous RBP levels and in an RBP’s normal spatiotemporal expression pattern.

BRAT and MEI-P26 are highly dosage-sensitive regulators: deviation from physiological levels is known to drive developmental defects in neuroblast lineages (*17–19, 24*), which can lead to lethality (*17, 25*). In contrast, flies carrying our Switch-TRIBE alleles exhibited no developmental defects, and single-cell analysis confirmed that neuroblast lineage composition was largely unaffected. This fidelity is reinforced by our heterozygous design, which maintains one wild-type allele. This configuration reduces the fraction of edited transcripts within the total RNA population, thereby minimizing potential impacts on RNA stability or translation mediated by A-to-G edits. Global gene expression remained largely unaltered in Switch-TRIBE cells, with the exception of two direct lncRNA targets in the BRAT dataset. Alterations in the expression level of one of these, *cherub*, is known not to impair neuroblast lineage development (*29*), consistent with the absence of phenotypic defects in our Switch-TRIBE experiments. Thus, Switch-TRIBE effectively maps interaction histories while preserving the native regulatory landscape.

Beyond preserving normal cellular physiology, rigorous target identification also requires the precise exclusion of technical noise (*13*). A major challenge in RNA-editing-based methods is the efficient removal of false positives arising from genomic variants (SNPs) and endogenous ADAR activity. TRIBE methods typically utilize matched genomic DNA to filter SNPs (*16*). However, this approach significantly increases sequencing costs and fails to account for background editing by endogenous ADARs, leading to potential false positives. Alternatively, using wild-type RNA as a control can measure endogenous editing (*14*) but risks overlooking strain-specific SNPs if the genetic background is not perfectly matched. Furthermore, computational filtering relies on comprehensive SNP databases, which remain unavailable for many non-model organisms. Switch-TRIBE addresses this limitation through a streamlined experimental design that utilizes the parental stocks as background controls. By sequencing the parents, we capture the exact repertoire of genetic variants and endogenous editing events inherited by the F1 experimental progeny. This approach effectively defines the ‘background noise’ without the need for additional genome sequencing or reliance on external databases. Thus, Switch-TRIBE offers a self-contained and universally applicable framework for target identification, readily adaptable to non-model organisms or diverse genetic backgrounds.

Leveraging this high-confidence detection framework, Switch-TRIBE has resolved the long-standing biological question regarding direct targets of BRAT in Drosophila neurogenesis, allowing us to categorize previous, genetically identified downstream effectors into direct and indirect targets. The method also validated the direct binding of MEI-P26 to its known downstream factor, *pros* (*24*). Furthermore, by identifying the targetomes of BRAT and MEI-P26, Switch-TRIBE has provided insights into the regulatory roles of these RBPs, identifying the mRNA encoding a novel regulator, 14-3-3ε, among the targets. Functional analysis further revealed that BRAT and MEI-P26 target genes encode core components of the metabolic and cytoplasmic translational machineries. Switch-TRIBE also characterized the consequences of co-binding of paralogous RBPs, BRAT and MEI-P26, in terms of target expression level and stability, extending our understanding of RBP regulatory outcomes. To place these findings in a broader context, we integrated our data with the recently published targetomes of IMP and SYP (*38*), two other crucial regulators of Drosophila neurogenesis (*39*). This comparison revealed extensive cross-regulation, where BRAT and MEI-P26 are targets of IMP and SYP, and *vice versa*. Furthermore, of 2,535 targets of IMP/SYP 1,861 (73.4%) were identified as targets of BRAT/MEI-P26 in our study (**Table S3**). These findings indicate the existence of a mutual regulatory network, suggesting that cell fate transitions are governed by highly interconnected layers of post-transcriptional control.

While established here in Drosophila, Switch-TRIBE employs a modular design to enable versatile profiling of context-dependent RBP functions across diverse systems. In *Drosophila*, distinct tissue-specific targetomes can be systematically interrogated by simply exchanging the Gal4 parental stock. This versatility extends to mammalian models and organoids via the universally available Cre-LoxP system. Beyond spatial control, this modularity can facilitate precise temporal regulation by using drug-inducible or optogenetic Cre drivers to capture dynamic RBP-RNA interactions. Furthermore, since Switch-TRIBE records interaction histories directly into the RNA sequence, it is inherently compatible with diverse sequencing platforms such as single-cell sequencing and long-read sequencing technologies, allowing for cell-type and isoform-specific resolution.

Switch-TRIBE also presents limitations. First, ADAR has an editing preference for adenosines within double-stranded RNA (*40*). This detection scope could be further improved in the future by utilizing different editing enzymes. Second, the method requires the genomic insertion of the LoxP-STOP-LoxP-ADARcd cassette via CRISPR/Cas9. Generating such knock-in strains can be time-consuming and may be challenging in species where genetic tools are limited. Third, fusing an RBP with an RNA editing enzyme could theoretically affect RBP-RNA interactions, and the A-to-G edits themselves might be toxic (*14, 16*). However, Switch-TRIBE minimizes these potential effects through its heterozygous design and endogenous expression levels.

In summary, Switch-TRIBE resolves the trade-off between detection sensitivity and biological fidelity. By capturing interaction histories at physiological levels, our approach provides a precise map of regulatory networks free from dosage-dependent artifacts. We anticipate that Switch-TRIBE will serve as a versatile framework to investigate the dynamic post-transcriptional mechanisms governing development and disease.

## MATERIALS AND METHODS

### Experimental model details

*Drosophila melanogaster* strains were cultivated under standard laboratory conditions at 25 °C. Strains included: *mei-P26*-*loxP-STOP-loxP-ADARcd* (MEI-P26 switch TRIBE, generated in this study); *brat*-*loxP-STOP-loxP-ADARcd* (BRAT switch TRIBE, generated in this study); UAS-Cre (BDSC#55800, BDSC#55801); *insc*-*Gal4* (BDSC#8751); *oc^otd^*/*FM7c*,*Tb*-*RFP* (BDSC#36337); *png^50^*/*FM6*; *Sp*/CyO, *tub-gal80;;act-gal4,UAS-GFP-nls/TM6B,Tb* (*41*); *14-3-3ε* RNAi (*42*) (BDSC#31497).

### Generation of switch-TRIBE stocks

CRISPR-mediated knock-in was performed by WellGenetics using modified methods based on Kondo and Ueda (*43*). Briefly, gRNA sequences targeting *brat* (CTATGTCCAACTGGCGCCAG[TGG]) or *mei-P26* (CGGACTAAAAATCACATCCG[AGG]) were cloned into U6-promoter plasmids. Donor plasmids were constructed in pUC57-Kan containing homology arms flanking the *loxP-3xStop-loxP-ADARcd-PBacDsRed* cassette, which encodes a V5-tagged ADARcd fusion protein and a 3xP3-DsRed marker flanked by *piggyBac* terminal repeats. *w^1118^* embryos were microinjected with a mixture of gRNA plasmids, donor plasmids, and an *hs-Cas9* plasmid. G0 adults were crossed, and F1 progeny were screened for the 3xP3-DsRed selection marker. Correct integration of the cassette was verified by genomic PCR using primer pairs targeting the 5’ and 3’ junctions. Specifically, primers were designed to anchor one end within the cassette and the other in the genomic region outside the homology arms, ensuring amplification only upon site-specific insertion. Upstream primers were BRAT-FWD TTCAGCAACGGAGGACTTCT; BRAT-REV ATTGGCAGGGGTACGTTCTT; MEI-P26-FWD GTTGCAAATTCGGTTCACTG; MEI-P26-REV TTGGCAGGGGTACGTTCTT. Downstream primers were BRAT-FWD TTTGACTCACGCGGTCGTTA; BRAT-REV TTCCTTGCGAACGAAGAAAT; MEI-P26-FWD TTTGACTCACGCGGTCGTTA; MEI-P26-REV TTATTGCCTGTACGCGACCT.

Genotyping confirmed specific amplification in heterozygous Switch-TRIBE samples and no amplification in negative controls. Validated transformants were balanced, and the *piggyBac* transposon containing the 3xP3-DsRed marker was subsequently excised by injecting a *piggyBac* transposase source. This excision retains a TTAA scar sequence designed to maintain the open reading frame. Final Switch-TRIBE stocks were validated by PCR for marker loss and established over balancer chromosomes.

### RBP-RNA immunoprecipitation

RBP-RNA immunoprecipitation (RIP) was performed using the anti-MEI-P26 D040 synthetic antigen-binding fragment (Fab), which was expressed and purified as previously described (*44*). Dissected *w^1118^* wandering L3 brains were homogenized in cold PBS, and the lysate was cleared by centrifugation at 20,000 g for 15 min at 4 °C. For each replicate, 40 μg of anti-FLAG agarose beads were pre-coated with 20 μg of the purified Fab in the presence of 1% BSA at 4 °C for >12 h. The cleared brain lysate was then incubated with the antibody-bead complex for 3 h at 4 °C. Following incubation, beads were washed four times with RIP wash buffer (150 mM KCl, 20 mM HEPES-KOH pH 7.5, 1 mM MgCl₂, 1 mM DTT, 1 mM AEBSF, 2 mM benzamidine, 2 μg/mL pepstatin, 2 μg/mL leupeptin, and 0.1% Triton X-100). Bound RNA was extracted directly from the beads using TRIzol reagent.

### RNA extraction and bulk RNA sequencing

Total RNA was extracted from dissected wandering L3 brains using TRIzol reagent (Invitrogen), following the manufacturer’s protocol, with the addition of 40 μg glycogen to facilitate precipitation. RNA integrity and concentration were assessed prior to library preparation.

Sequencing libraries were constructed using NEBNext Ultra RNA Library Prep Kit for Illumina (NEB, USA, Catalog #: E7530L) and sequenced on an Illumina NovaSeq 6000 platform (Novogene) in paired-end 150-bp (PE150) mode.

### Single-cell RNA sequencing

Single-cell Switch-TRIBE libraries were constructed using the Singleron platform. Dissected wandering L3 brains were preserved in sCelLive Tissue Preservation Solution (Singleron Biotechnologies) and dissociated into single-cell suspensions using a PythoN Automated Tissue Dissociator with the sCelLive Tissue Dissociation Mix. Cell viability was assessed via trypan blue staining. Single-cell suspensions were adjusted to 1 × 10^5^ cells/mL in PBS and loaded onto microfluidic chips. For both MEI-P26-Switch-TRIBE and BRAT-Switch-TRIBE samples, around 30,000 cells were loaded per genotype. scRNA-seq libraries were constructed using the GEXSCOPE Single-Cell RNA Library Kit on a Matrix Automated Single-Cell Processing System. Libraries were pooled and sequenced on an Illumina NovaSeq 6000 platform in PE150 mode.

### Antibodies and Immunostaining

Dissected L3 brains were fixed in 4% formaldehyde for 20 min at room temperature (RT) and washed three times in PBS containing 0.3% Triton X-100 (PBST). Tissues were blocked with 1% bovine serum albumin (BSA) in PBST for 1 h at room temperature, followed by incubation with primary antibodies overnight at 4 °C. After washing (3 × 10 min in PBST), samples were incubated with secondary antibodies for 2 h at RT. Tissues were washed again (3 × 10 min in PBST), mounted in antifade medium (1% DABCO, 90% glycerol in PBS), and imaged using a Leica DMi8 TCS SP8 confocal microscope.

The following primary antibodies were used: mouse anti-V5 (Abcam, ab27671; 1:500), rat anti-ELAV (Developmental Studies Hybridoma Bank, 7E8A10; 1:50), rabbit anti-MEI-P26 (*45*) (1:500), rat anti-BRAT (*18*) (1:500), and rabbit anti-BRAT (*46*) (1:500). Donkey and goat secondary antibodies conjugated to Alexa Fluor 488, 555 and 647 were used (1:300, ThermoFisher Scientific).

### Identification of bulk Switch-TRIBE A-to-G edits

Switch-TRIBE editing sites were identified following published HyperTRIBE protocols (*14, 16*). with modifications. For both BRAT and MEI-P26 experiments, three biological replicates were collected for each sample type: (1) Switch-TRIBE (experimental group); (2) *insc-Gal4* only (control); and (3) *UAS-Cre* cassette only (control). Sequencing reads were aligned to the *Drosophila melanogaster* reference genome (r6.54) using *STAR* (v2.7.10a) (*47*). PCR duplicates and low-quality reads were removed using *samtools* (v1.13) (*48*) and *Picard MarkDuplicates* (v2.27.1). The processed BAM files were converted into matrix format and imported into a MySQL (MariaDB) database for downstream analysis.

In each Switch-TRIBE replicate, candidate A-to-G editing sites were identified by selecting A-dominant loci (adenosine accounting for more than 50% of total reads) with non-zero guanosine (G) counts. To ensure reproducibility, candidate sites were required to be present in at least two out of three Switch-TRIBE replicates. To eliminate false positives arising from SNPs, we excluded any locus displaying G-dominance in any of the six control replicates. For the remaining candidates, A and G counts from Switch-TRIBE replicates were aggregated and compared against each of the two control groups independently using Chi-square tests. Sites were retained if they met one of the following criteria: (1) significant differences (Bonferroni-adjusted *P* < 0.05) were observed in comparisons with both control groups; or (2) significant differences were observed in one comparison, provided that the other control group had an accumulated read depth lower than 30. To capture A-to-G editing events on transcripts originating from the genomic negative strand, the same pipeline was applied to identify T-to-C mismatches.

### Single-cell RNA sequencing data processing and cell type annotation

Single-cell sequencing data were processed using *STARsolo* (v2.7.11b) with the Drosophila reference genome (r6.54) and the cell barcode whitelist provided by Singleron (https://github.com/singleron-RD/CeleScope). The output matrix was analyzed using the R package *Seurat* (v5.3.0) (*49*). Low-quality cells (<200 unique features or >20% mitochondrial RNA) were removed, retaining 20,553 cells for BRAT and 21,393 cells for MEI-P26 Switch-TRIBE datasets. Gene expression matrices were normalized using NormalizeData and scaled using ScaleData. The top 2,000 variable genes were identified via FindVariableFeatures. Dimensionality reduction and clustering were performed using the top 15 principal components, the Leiden algorithm (resolution = 1.0), and visualized using t-SNE.

Cell types were annotated based on a curated panel of 27 marker genes derived from previous studies (*50, 51*). Clusters were defined as follows: Neuroblasts (NB): *dpn*+, *CycE*+, *lncRNA:CR31386*+; Intermediate Neural Progenitors (INP): *dpn*+, *CycE*+, *erm*+, *ham*+, *ytr*+, *D*+, *ase*+, *mira*+; Ganglion Mother Cells (GMCs) were subdivided into Type I lineage (GMCT1: *dap*+, *tap*+, *insb*+; *erm*-, *ham*-, *ytr*-, *D*-) and Type II lineage (GMCT2: *dap*+, *tap*+, *insb*+, *erm*+, *ham*+, *ytr*+, *D*+); Neurons: *nSyb*+, *CadN*+, *elav*+; and Glia: *repo*+. Type II neuroblasts were not identified, likely due to their rarity. Downstream analysis was restricted to cells originating from Type I and Type II neuroblast lineages in the central brain, selected based on annotation and cluster topology (**Figs. S3C, S4C**).

### Identification of single-cell Switch-TRIBE A-to-G edits

To identify editing events in single-cell RNA-seq data, we implemented a targeted approach leveraging the high-confidence sites defined in bulk Switch-TRIBE. First, we identified Switch-TRIBE-positive single cells by detecting R2 sequences harboring the Cre-LoxP-recombination scar using seqkit. Due to the sparse coverage of scRNA-seq, we utilized the bulk data to define a reference set of high-confidence A-to-G editing sites, applying stringent criteria: (1) A-dominant in all six control replicates with zero G counts; and (2) more than three G counts and more than 2.5% editing frequency in all three bulk Switch-TRIBE replicates.

Given these stringent criteria, we operated under the assumption that any scRNA-seq R2 read (harboring the sequence of the actual RNA fragment) containing these specific A-to-G edits represents Switch-TRIBE activity. We subsequently extracted cellular barcodes from the paired R1 reads (harboring the cellular barcodes and UMIs) corresponding to these R2 sequences and utilized them to label Switch-TRIBE-positive cells in Seurat. Following cell-type annotation, R2 reads from Switch-TRIBE-positive cells were aggregated by cell type to generate pseudobulk samples. These pseudobulk reads were processed using STAR and the MySQL pipeline described above. To rigorously distinguish genuine editing events from genomic variants or sequencing errors, we validated potential edits in the pseudobulk samples by performing Chi-square tests against each of the six bulk control replicates individually. Only sites maintaining significance (Bonferroni-adjusted *P* < 0.05) were retained as single-cell editing events.

### Distance distribution analysis

The genomic distances between BRAT and MEI-P26 editing sites were calculated based on their mapped positions within the longest isoform of each target transcript. Sites located outside the exonic regions of the longest isoforms were excluded from the analysis.

### Motif enrichment analysis

Motif occurrence (NOM) and sum of accessibility (SOM) were analyzed using the custom developed MFRA (Motif and Flanking Region Accessibility) pipeline (available at github.com/LipshitzLabToronto). Briefly, target and non-target transcript sets were defined based on the Switch-TRIBE data, and RNA secondary structure accessibility was predicted using *RNAplfold* (ViennaRNA Package v2.5.1) (*52*) with the parameters -W 80 -L 40 -u 10. Target and non-target transcripts, together with their corresponding *RNAplfold* accessibility matrices, were analyzed using the MFRA algorithm to identify the occurrence of the predefined motif with the highest predicted accessibility in each transcript and to extract accessibility scores for its 5′ and 3′ flanking regions (4-8 nucleotides). Mean and median accessibility scores were calculated separately for target and non-target transcript sets, and statistical significance was assessed using a two-sided Mann-Whitney U test (*P* < 0.05).

### Edit-to-motif Distance analysis

To assess the spatial proximity of editing events to binding motifs, we calculated the distance from each editing site to its nearest identified motif based on genomic coordinates. The consensus binding motifs were defined as UGUUD for BRAT and a combination of UUUAUC, UUUACA, UUUAAU, UUUCUA, UUUUAC and UUUUUU for MEI-P26, based on previous studies (*21, 22*). Control motifs were selected from sequences exhibiting the lowest binding affinity in published RNAcompete datasets (*53*): AGUVA for BRAT and a combination of GUGGAG, AGGUGG, GGGUGA, AAGUAG, AGUGAG and UGUGAG for MEI-P26.

### Phenotypic analysis of *14-3-3ε*

Mosaic neuroblast lineages depleted of *14-3-3ε* were generated using the Illuminati method as previously described (*24, 41*). Briefly, Illuminati driver stocks were crossed to *UAS-14-3-3ε-RNAi* lines to generate Illuminati F1 progeny. Wandering third-instar larval brains were dissected, fixed, and imaged using a Leica DMi8 TCS SP8 confocal microscope. Lineage size was quantified by measuring the total volume of GFP-positive clones using LAS X software. The genotype of the control was *tub-gal80;;act-gal4,UAS-GFP-nls/TM6B*.

### RIP-Seq target identification

Raw sequencing reads from MEI-P26 RIP-seq and input controls (n=3 biological replicates) were processed using *Trim Galore* (v0.6.10) for quality control and *SortMeRNA* (v4.3.6) to remove rRNA. Transcript abundance was quantified directly from the filtered reads using *Salmon* (v1.10.2) (*54*) against an index derived from the *Drosophila melanogaster* reference transcriptome. Differential gene expression analysis was performed using *DESeq2* (*55*), with significance defined by adjusted *P* values.

### Gene Ontology (GO) enrichment analysis

Functional enrichment analysis was performed using g:Profiler (*56*). The lists of high-confidence target genes were tested for enrichment of Gene Ontology (GO) terms against the *Drosophila melanogaster* database. Statistical significance was determined using the g:SCS threshold method with a cutoff of adjusted *P* < 0.05.

### Statistical tests of Euler diagram overlaps

The significance in Euler diagrams (**Fig. 5C, 5J**) was determined by the hypergeometric test. In **Fig. 5C**, the total number of genes expressed in either early embryo or L3 brain was 12,610; while in **Fig. 5J** the total number of candidate A sites in the L3 brain transcriptome was 9,270,810.

## Acknowledgements Funding

This research was supported by grants from the Canadian Institutes of Health Research to H.D.L. (PJT-190124) and the International School of Medicine, Zhejiang University to X.Y. (Startup Fund Q23019). Y.H. was supported in part by University of Toronto Open Fellowships.

## Data and materials availability

The Switch-TRIBE Drosophila lines are available upon request from Howard Lipshitz or Xiaohang Yang. The datasets for RNA sequencing presented in this study are available in the Gene Expression Omnibus (GEO) repository under accession numbers GSE336850 (MEI-P26 brain RIP-seq), GSE336851 (Switch-TRIBE BRAT and MEI-P26 bulk brain RNA-seq), and GSE336852 (Switch-TRIBE BRAT and MEI-P26 brain single cell RNA-seq). The MFRA code is available at github.com/LipshitzLabToronto. All other data are present in the paper or the Supplementary Materials.

## Supplementary Materials

**Fig. S1.**
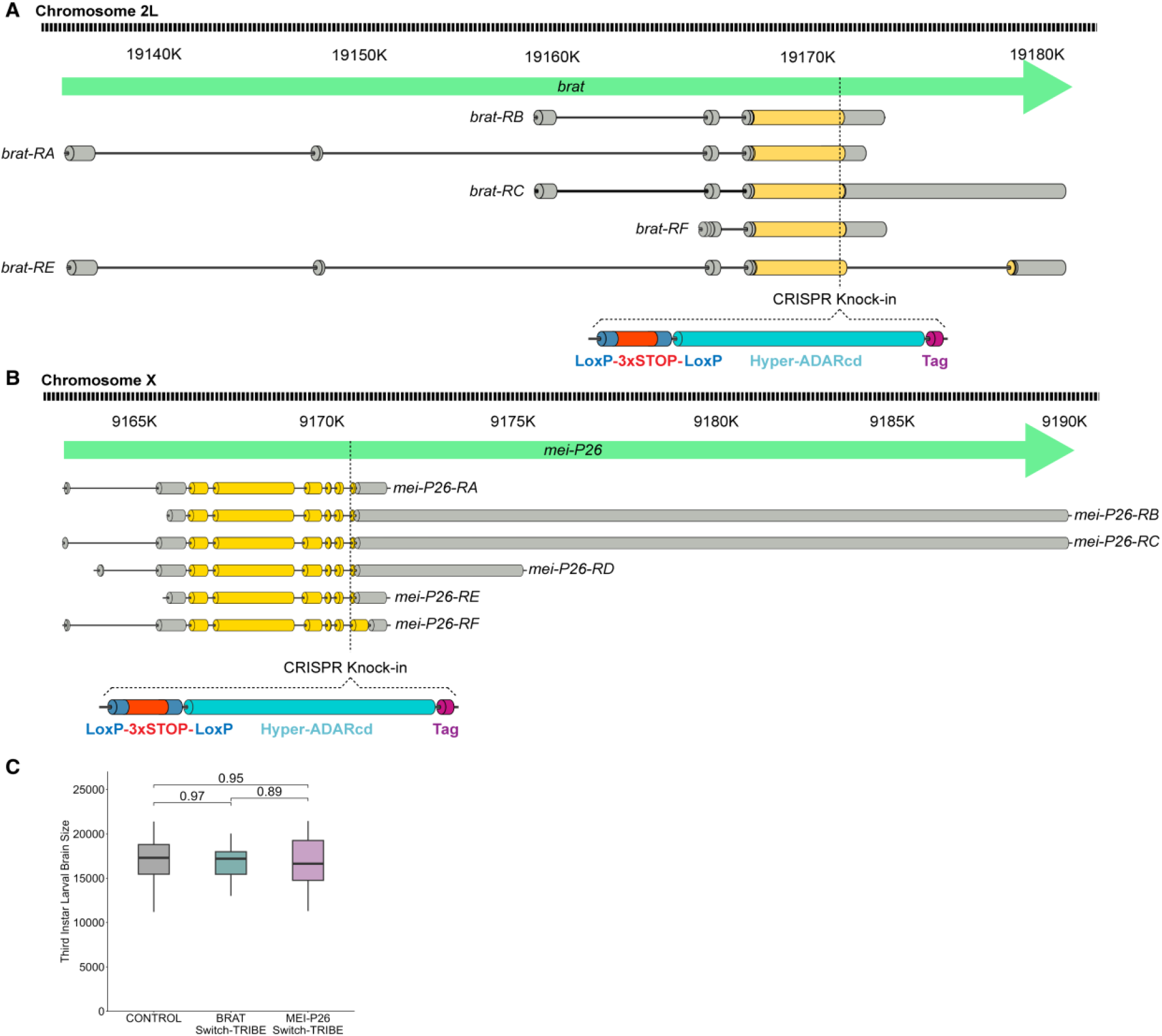
Generation of Switch-TRIBE CRISPR knock-in stocks. **(A)** Diagram of BRAT Switch-TRIBE CRISPR knock-in construct. The Switch-TRIBE cassette (LoxP-3xSTOP-LoxP-ADARcd-V5-His) was inserted into the 3’ of the *brat/CG10719* coding intron (Chromosome 2L, 37C1-37C6) of the donor *w^1118^* Drosophila stock via CRISPR knock-in techniques, creating a conditionally expressed ADAR fusion domain on the C-terminus of most BRAT isoforms. Exact insertion site and sequence can be found in **Figure S2**. **(B)** Diagram of MEI-P26 Switch-TRIBE CRISPR knock-in construct. The Switch-TRIBE cassette (LoxP-3xSTOP-LoxP-ADARcd-V5-His) was inserted into the 3′ of the *mei-P26/CG12218* coding intron (Chromosome X, 8C15-8C17) of the donor *w^1118^* Drosophila stock via CRISPR knock-in techniques, creating a conditionally expressed ADAR fusion domain on the C-terminus of most MEI-P26 isoforms. Exact insertion site and sequence can be found in **Figure S2**. **(C)** Quantification of third-instar larval brain sizes of BRAT-Switch-TRIBE, MEI-P26-Switch-TRIBE and control flies. (Bonferroni-corrected *P* values are from the Wilcoxon rank-sum test, n = 9, 16, 12, respectively.)

**Fig. S2:**
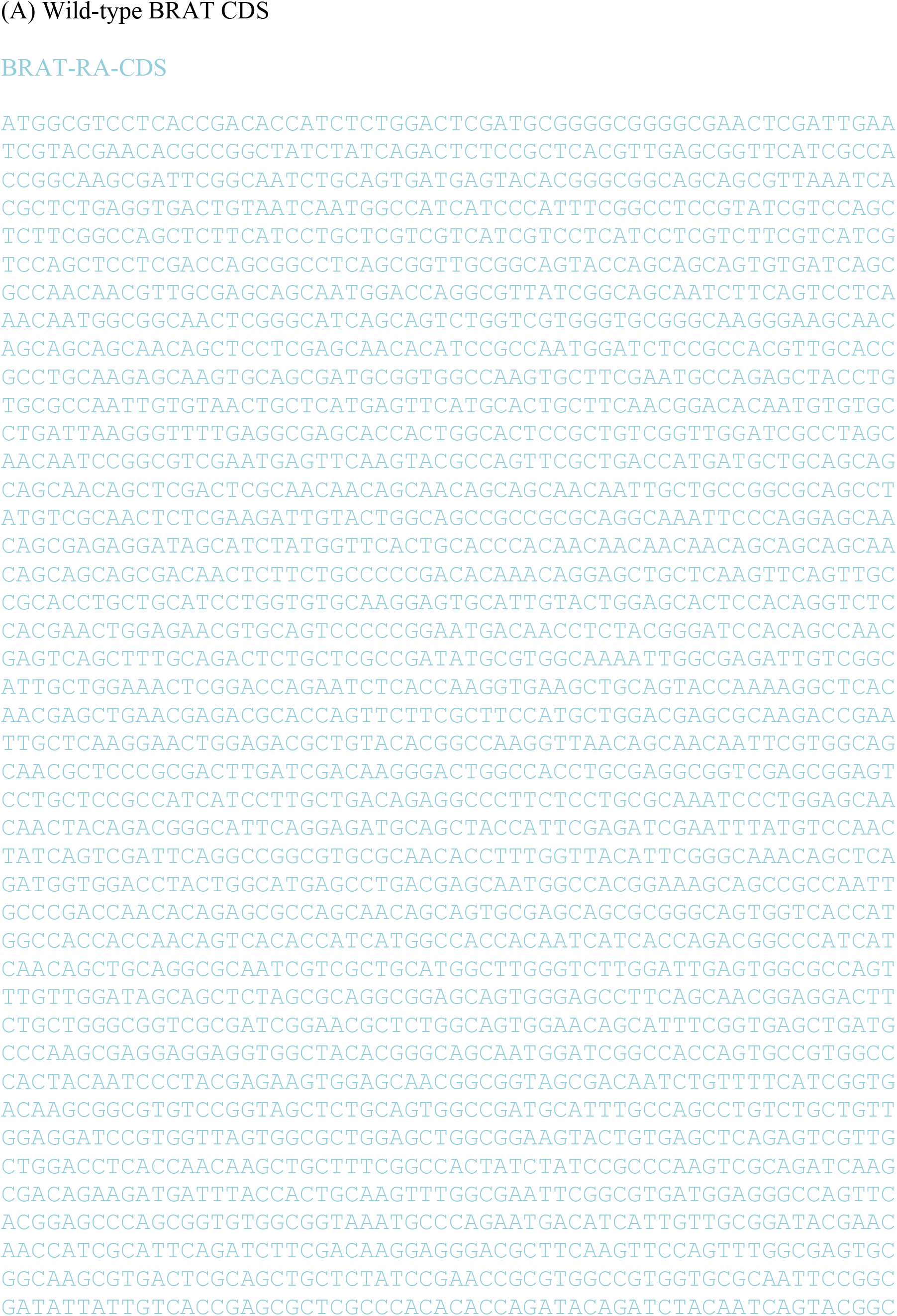
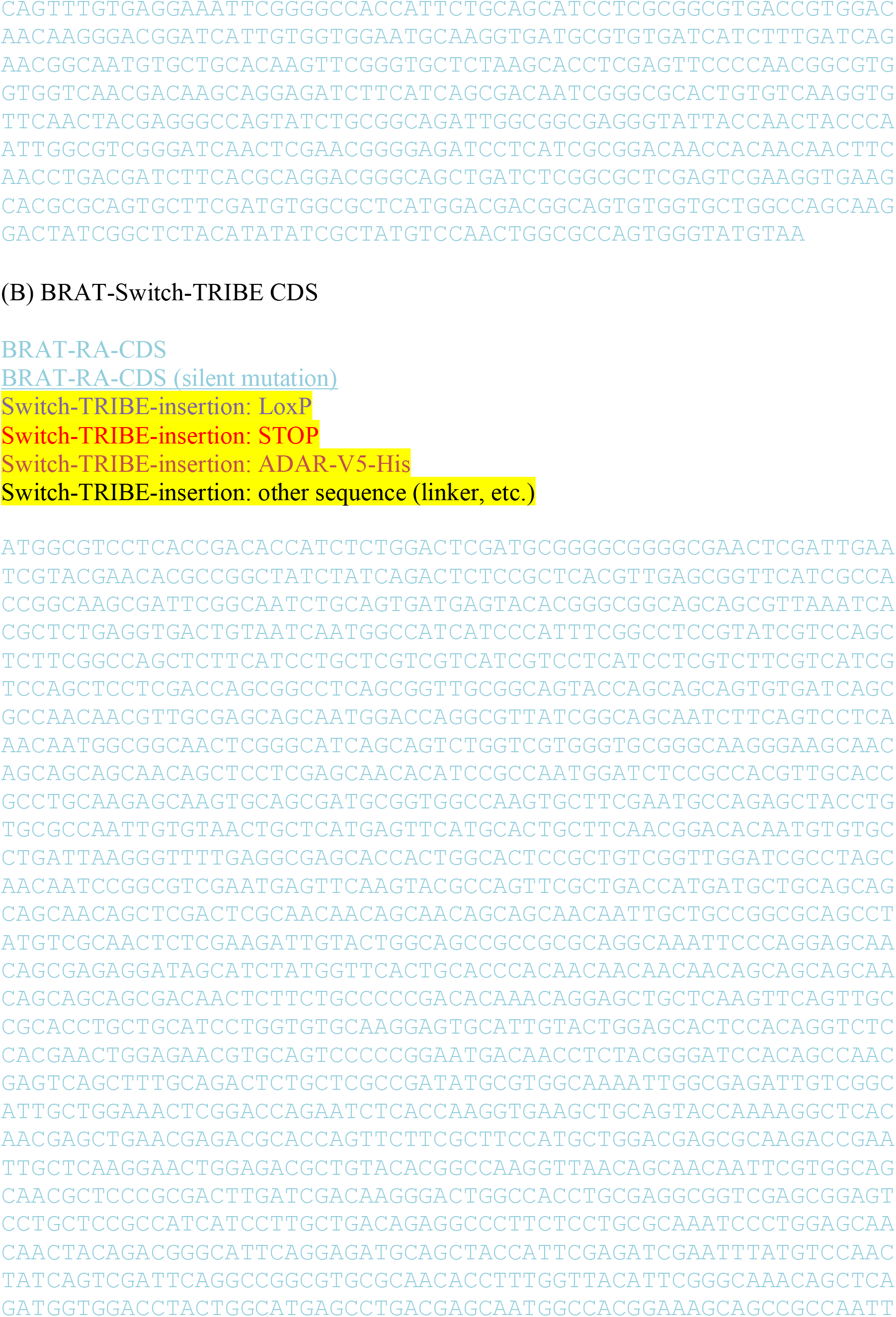
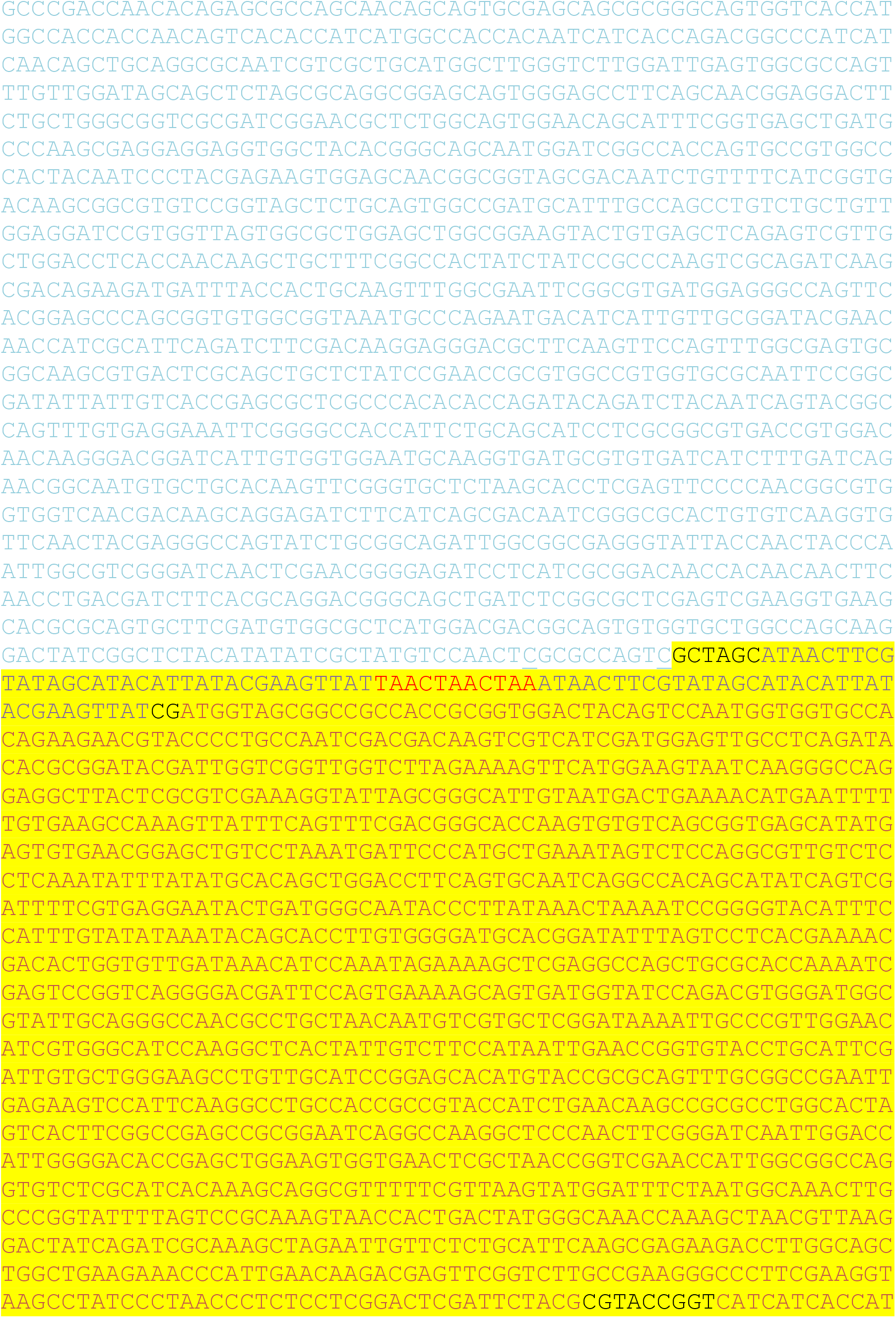
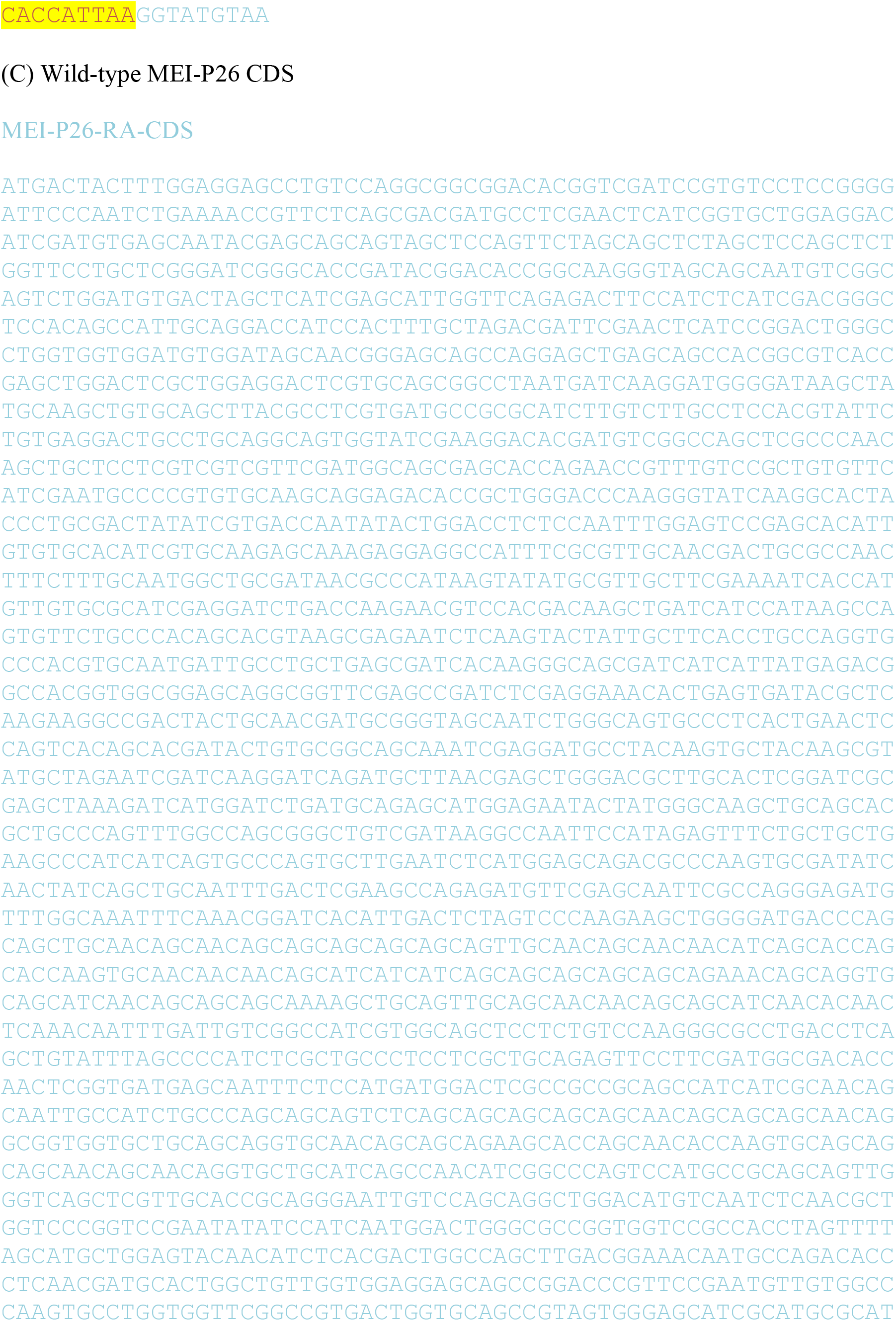
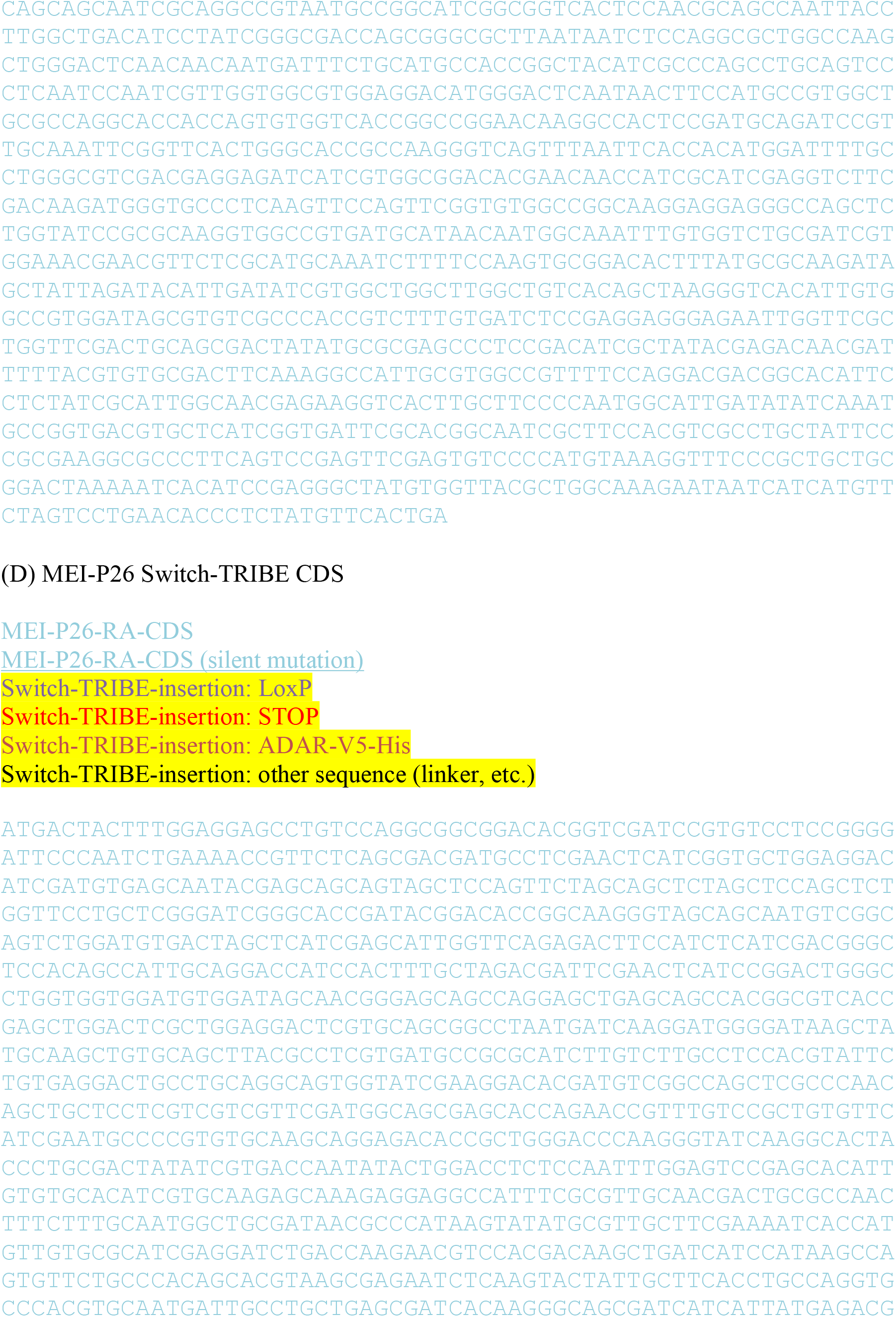
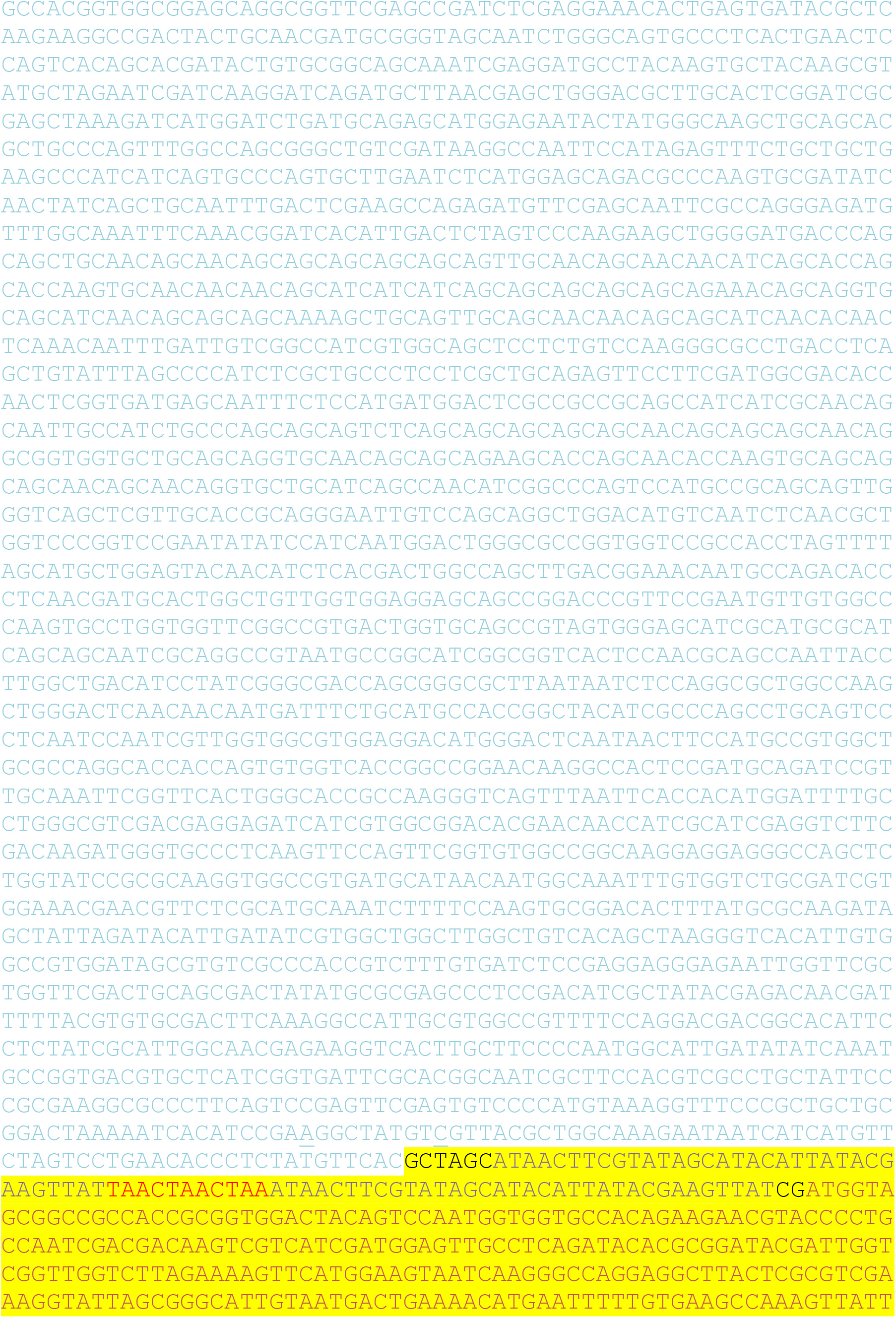
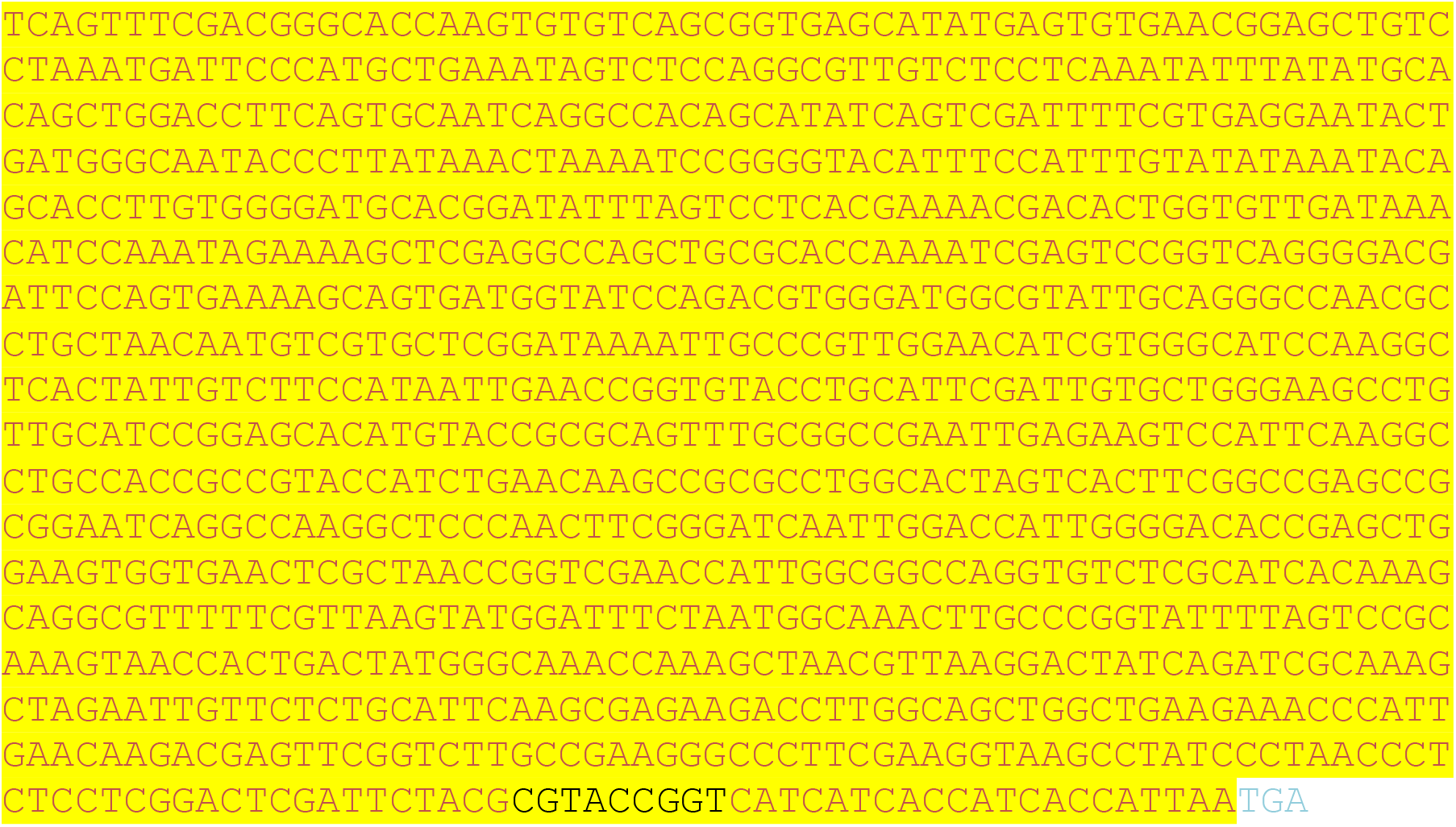
Sequence of unedited and CRISPR-edited *brat* and *mei-P26* loci. **(A)** Unedited *brat* coding sequence (CDS). **(B)** CRISPR-edited Switch-TRIBE *brat* CDS. **(C)** Unedited *mei-P26* CDS. **(D)** CRISPR-edited Switch-TRIBE *mei-P26* CDS.

**Fig. S3.**
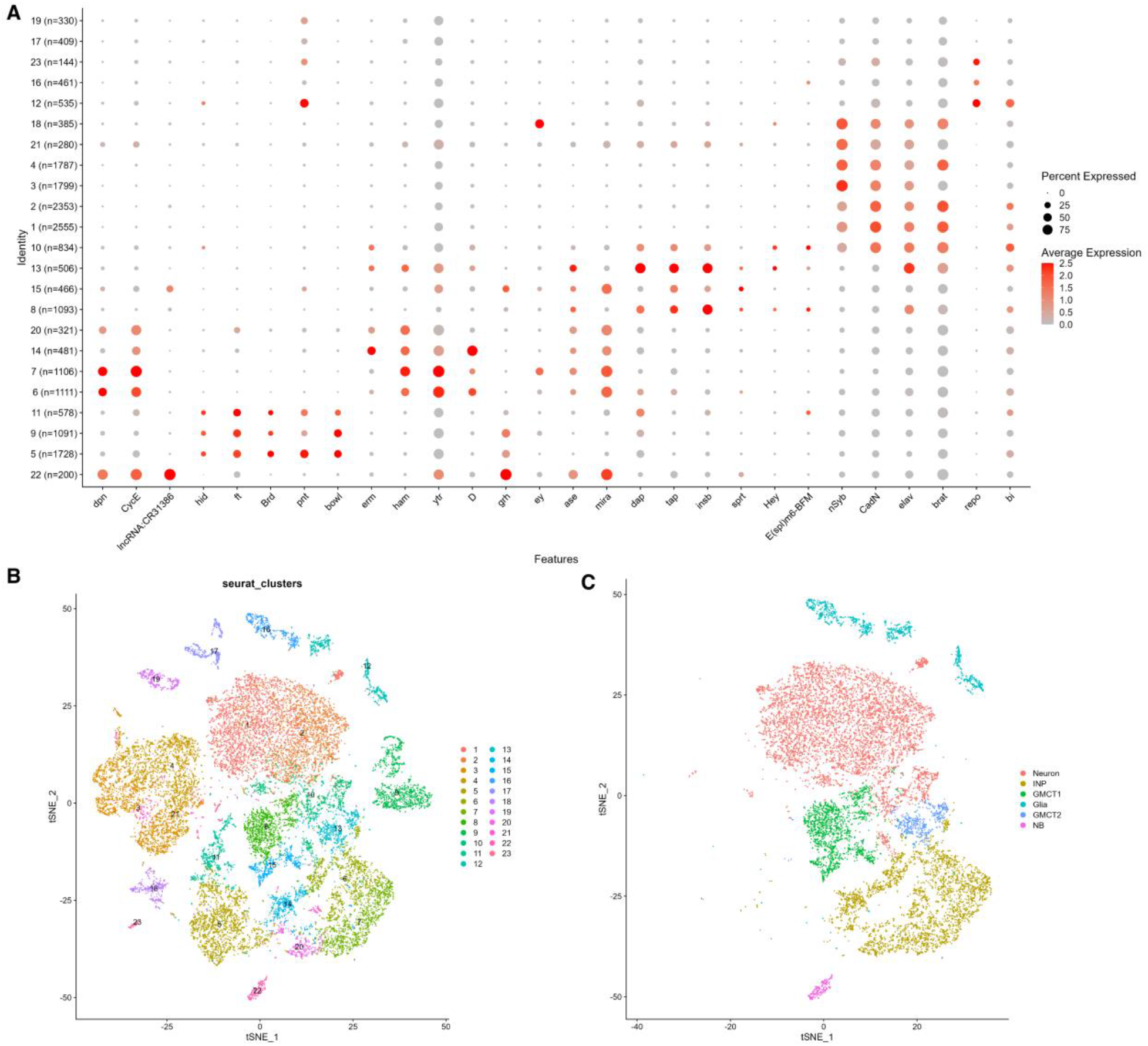
Cell type clustering in BRAT Switch-TRIBE samples. **(A)** BRAT single-cell Switch-TRIBE data was clustered into 23 cell clusters. Dot plot shows expression of cell-type marker genes in each cell cluster. **(B)** tSNE of BRAT Switch-TRIBE samples, colors represent clusters. **(C)** tSNE of BRAT Switch-TRIBE samples, colors represent six major cell types that are derived from Type I or Type II neuroblast lineages: NB (cluster 22), INP (cluster 6,7,14,20), GMCT1 (cluster 8,15), GMCT2 (cluster 13), Neuron (cluster 1,2,10) and Glia (cluster 12,16). Cells that are not clustered into these six major cell types were excluded from following analysis.

**Fig. S4.**
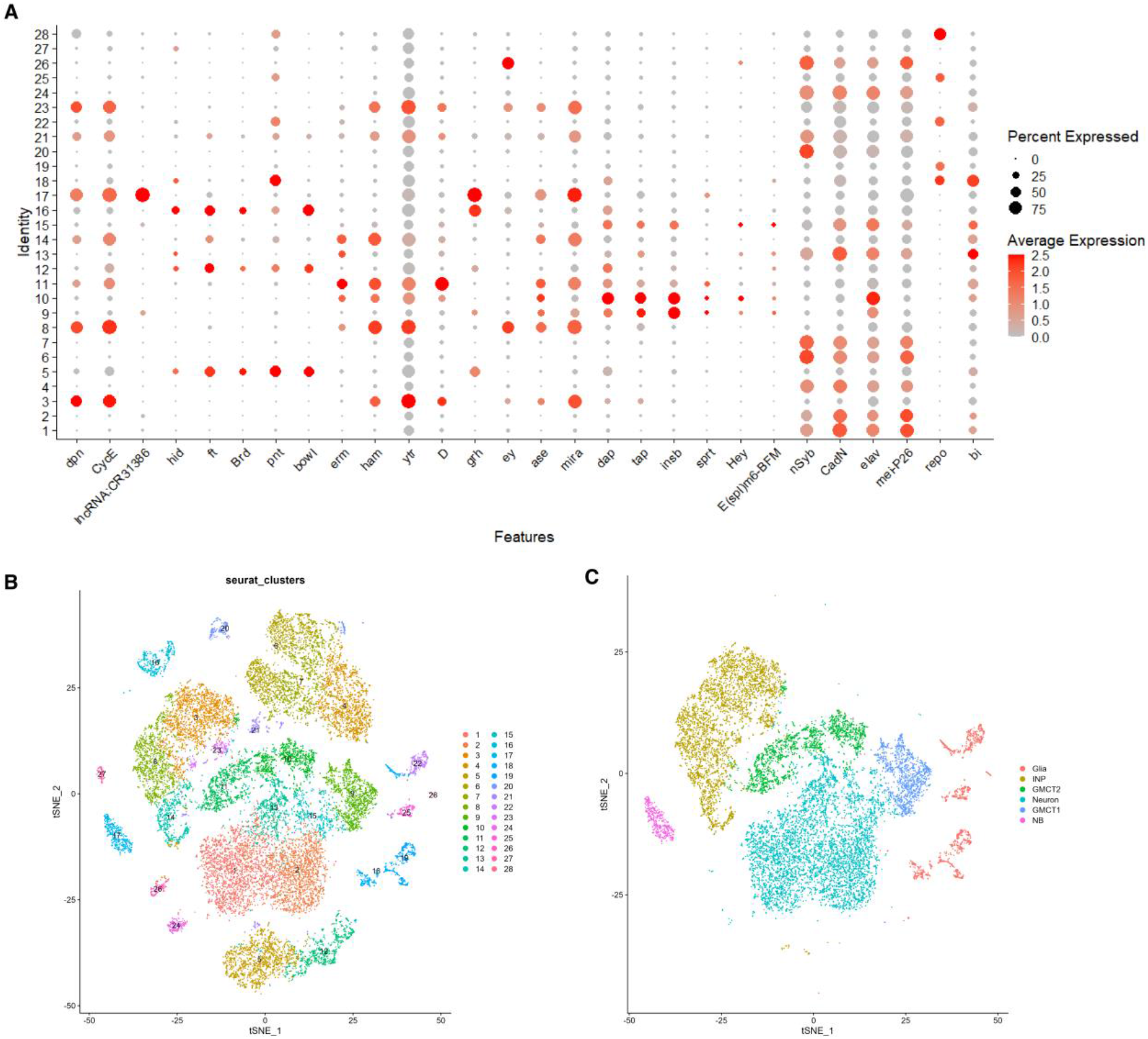
Cell type clustering in MEI-P26 Switch-TRIBE samples. **(A)** MEI-P26 single-cell Switch-TRIBE data was clustered into 28 cell clusters. Dot plot shows expression of cell-type marker genes in each cell clusters. **(B)** tSNE of MEI-P26 Switch-TRIBE samples, colors represent clusters. **(C)** tSNE of MEI-P26 Switch-TRIBE samples, colors represent six major cell types that are derived from Type I or Type II neuroblast lineages: NB (cluster 17), INP (cluster 3,8,14,23), GMCT1 (cluster 9), GMCT2 (cluster 10,11), Neuron (cluster 1,2,13,15) and Glia (cluster 18,19,22,25,28). Cells that are not clustered into these six major cell types were excluded from following analysis.

**Fig. S5.**
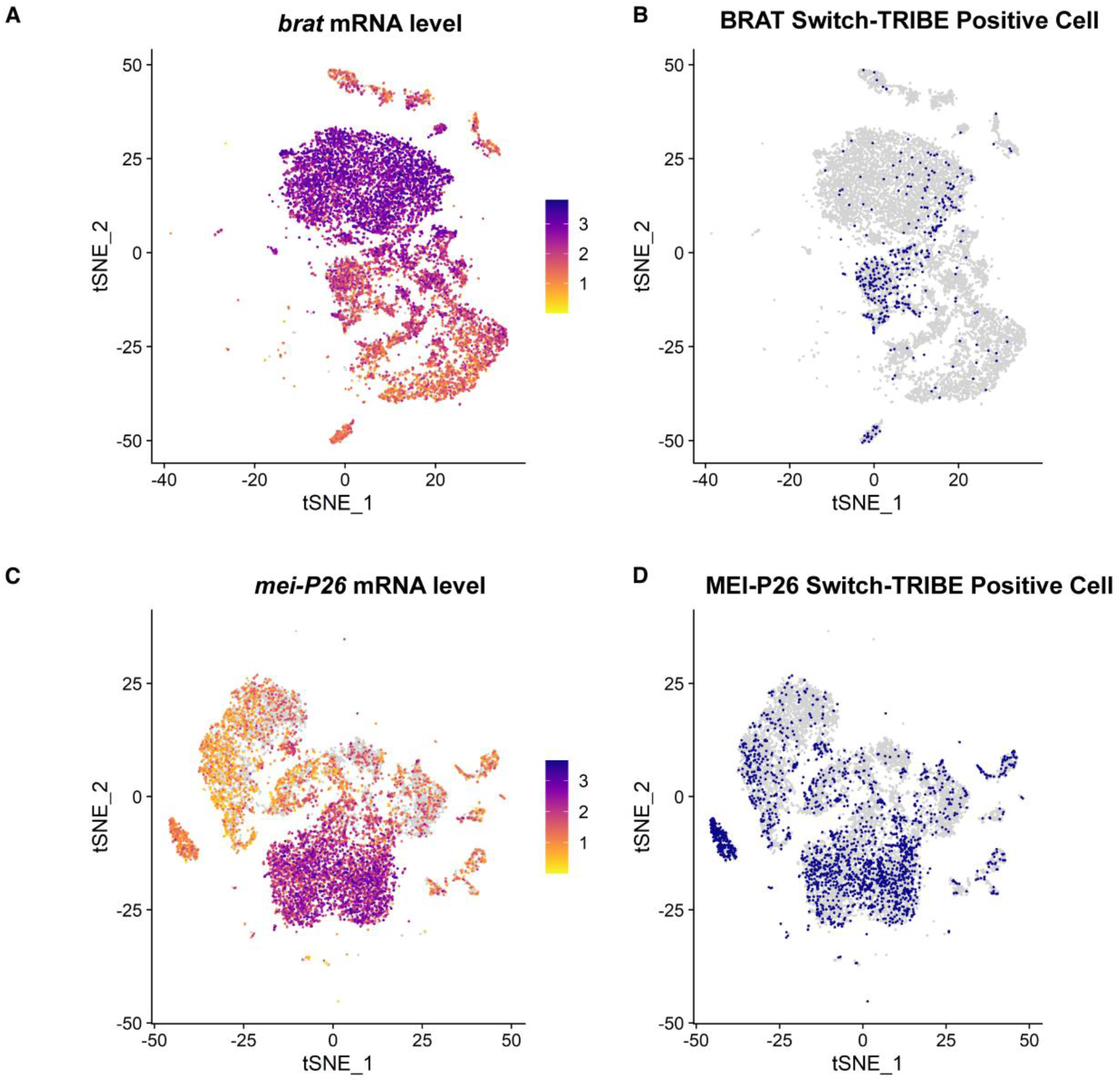
Visualization of Switch-TRIBE positive single cells. **(A)** tSNE of BRAT Switch-TRIBE samples (only 6 major cell types included). Each dot represents a single cell. *brat* mRNA levels in each individual cell are represented in color. **(B)** tSNE of BRAT Switch-TRIBE samples (only 6 major cell types included). BRAT Switch-TRIBE positive single cells are shown in blue, while Switch-TRIBE negative single cells are shown in grey. **(C)** tSNE of MEI-P26 Switch-TRIBE samples (only 6 major cell types included). *mei-P26* mRNA levels in each individual cell are represented in color. **(D)** tSNE of MEI-P26 Switch-TRIBE samples (only 6 major cell types included). MEI-P26 Switch-TRIBE positive single cells are shown in blue, while Switch-TRIBE negative single cells are shown in grey.

**Fig. S6.**
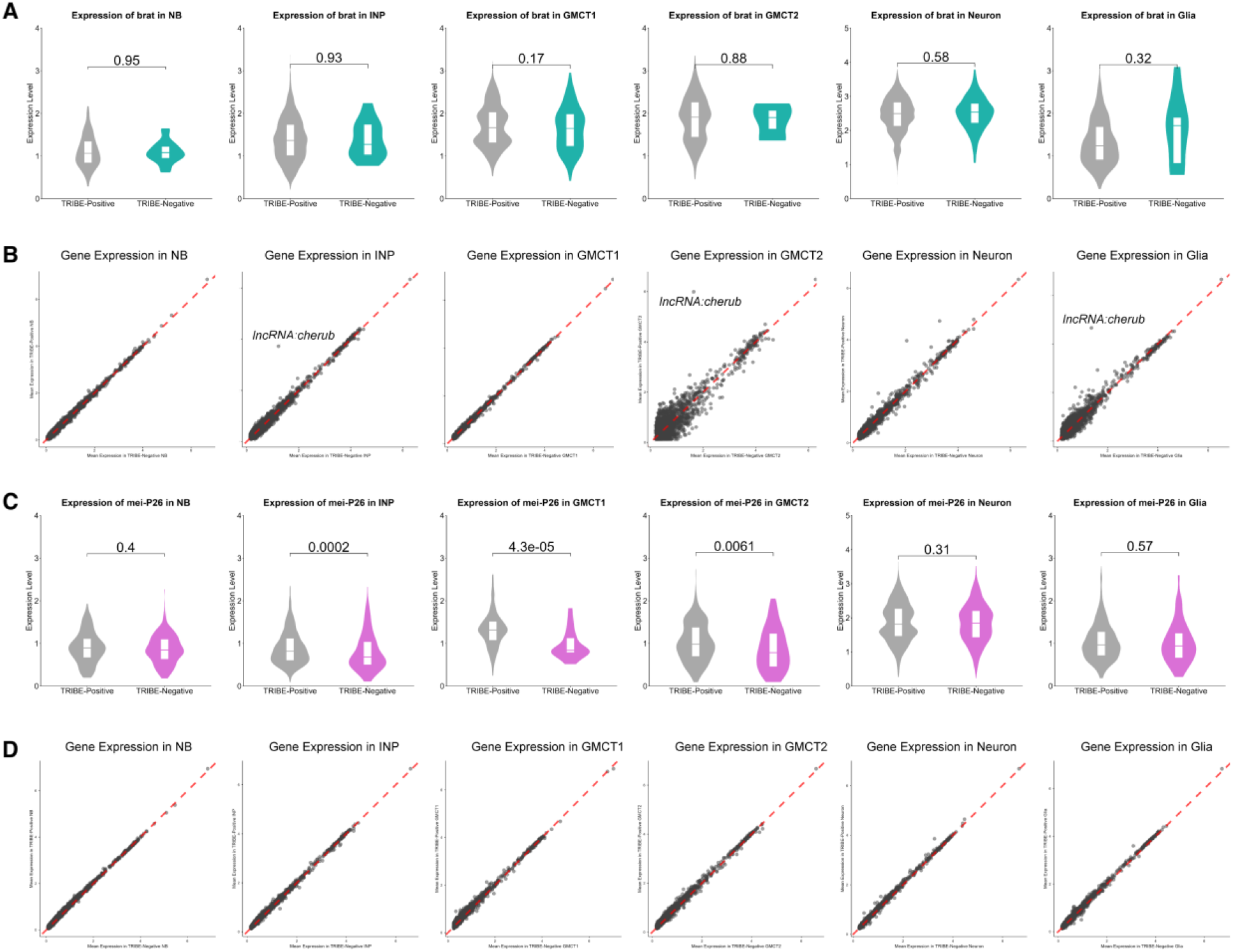
Switch-TRIBE has little effect on local transcriptomes. **(A)** The expression level of *brat* mRNA remains largely unchanged in Switch-TRIBE positive cells compared to Switch-TRIBE negative cells. Left to right: six different cell types. **(B)** The levels of expressed genes in BRAT Switch-TRIBE positive cells remain mostly unchanged, with the exception of *lncRNA*:cherub. Left to right: six different cell types. **(C)** The expression level of *mei-P26* mRNA remains largely unchanged in Switch-TRIBE positive cells in MEI-P26’s predominant cell types (NBs and neurons) compared to Switch-TRIBE negative cells. Left to right: six different cell types. **(D)** The levels of expressed genes in MEI-P26 Switch-TRIBE positive cells remain mostly unchanged. Left to right: six different cell types.

**Fig. S7.**
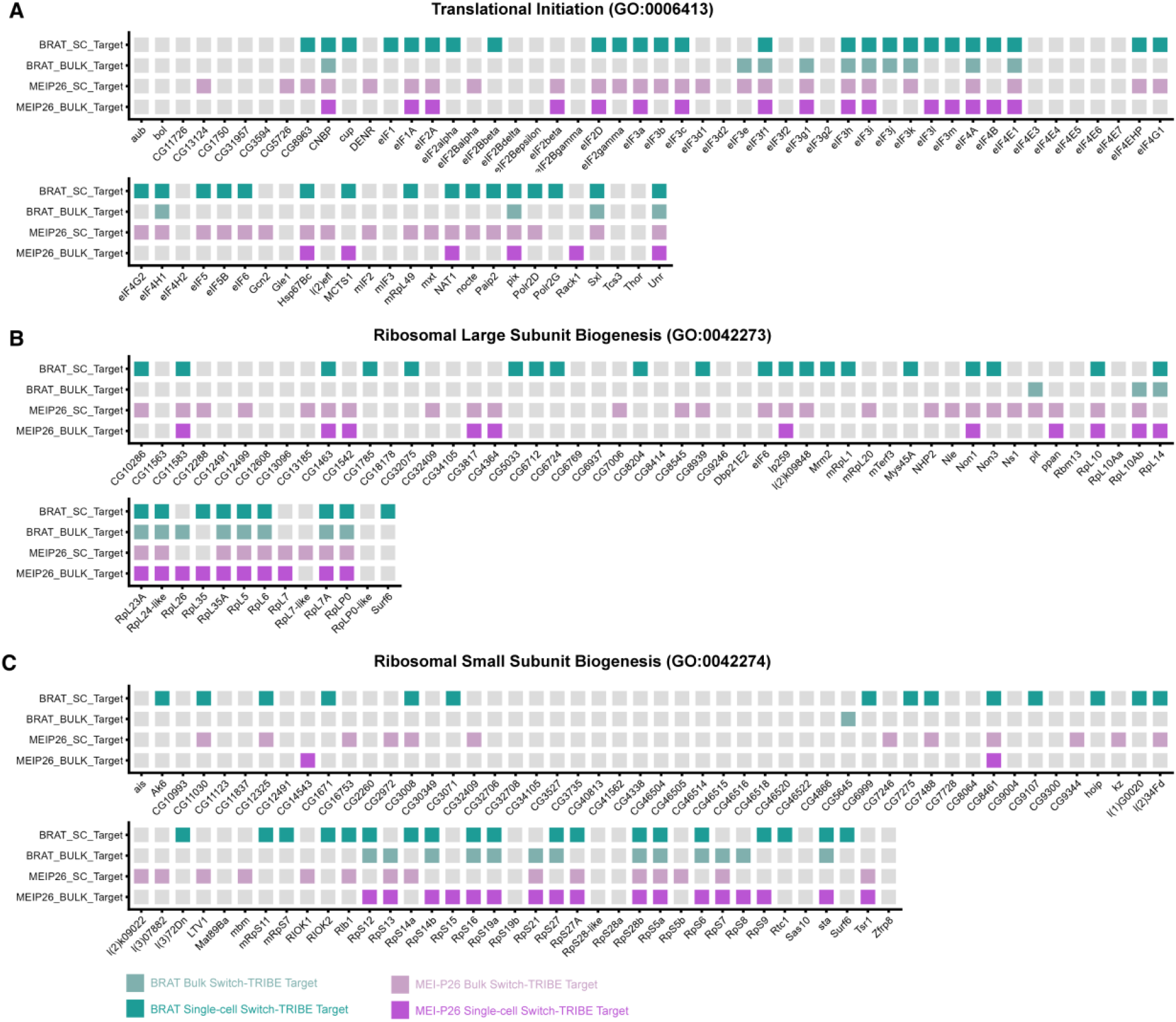
Map of BRAT and MEI-P26 direct targets within enriched GO terms. **(A)** Schematic of the GO term ‘Translational Initiation’ (GO:0006413); genes identified as direct BRAT and/or MEI-P26 targets are highlighted. BRAT and MEI-P26 directly target 54 out of 76 genes. **(B)** Schematic of the GO term ‘Ribosomal Large Subunit Biogenesis’ (GO:0042273); genes identified as direct BRAT and/or MEI-P26 targets are highlighted. BRAT and MEI-P26 directly target 48 out of 63 genes. **(C)** Schematic of the GO term ‘Ribosomal Small Subunit Biogenesis’ (GO:0042274); genes identified as direct BRAT and/or MEI-P26 targets are highlighted. BRAT and MEI-P26 directly target 53 out of 87 genes.

---

**Table S1. Targetomes of BRAT and MEI-P26 identified by bulk L3 brain and single-cell L3 brain Switch-TRIBE.** The start and end coordinates are given for each edited gene along with the coordinate(s) of the edited site(s). For single-cell Switch-TRIBE, the cell type(s) in which the edit(s) were detected is listed. P_ctr_1 is the multi-test-corrected *P* value of chi-squared test between Switch-TRIBE and the Gal4-only-control, while P_ctr_2 is the multi-test-corrected *P* value for chi-squared test between Switch-TRIBE and the cassette-only-control. The table is provided as a separate file.

**Table S2. Gene Ontology (GO) term analysis of BRAT and MEI-P26 targetomes identified by bulk L3 brain Switch-TRIBE.** For each GO term, the adjusted *P*-value, negative log10 of the adjusted *P* value, term size, query size, intersection, effective domain size (the total pool of genes considered for the hypergeometric test) are listed. g:Profiler automatically ‘highlighted’ a subset of the terms. The table is provided as a separate file.

**Table S3.** Overlap of BRAT and MEI-P26 targetomes identified by bulk and single-cell L3 Switch-TRIBE with published targets of IMP and SYP in brain as identified (*38*). The table is provided as a separate file.

